# Rapid isolation of potent SARS-CoV-2 neutralizing antibodies and protection in a small animal model

**DOI:** 10.1101/2020.05.11.088674

**Authors:** Thomas F. Rogers, Fangzhu Zhao, Deli Huang, Nathan Beutler, Alison Burns, Wan-ting He, Oliver Limbo, Chloe Smith, Ge Song, Jordan Woehl, Linlin Yang, Robert K. Abbott, Sean Callaghan, Elijah Garcia, Jonathan Hurtado, Mara Parren, Linghang Peng, James Ricketts, Michael J. Ricciardi, Stephen A. Rawlings, Davey M. Smith, David Nemazee, John R. Teijaro, James E. Voss, Raiees Andrabi, Bryan Briney, Elise Landais, Devin Sok, Joseph G. Jardine, Dennis R. Burton

## Abstract

The development of countermeasures to prevent and treat COVID-19 is a global health priority. In under 7 weeks, we enrolled a cohort of SARS-CoV-2-recovered participants, developed neutralization assays to interrogate serum and monoclonal antibody responses, adapted our high throughput antibody isolation, production and characterization pipeline to rapidly screen over 1000 antigen-specific antibodies, and established an animal model to test protection. We report multiple highly potent neutralizing antibodies (nAbs) and show that passive transfer of a nAb provides protection against high-dose SARS-CoV-2 challenge in Syrian hamsters. The study suggests a role for nAbs in prophylaxis, and potentially therapy, of COVID-19. The nAbs define protective epitopes to guide vaccine design.

## Introduction

The novel coronavirus disease (COVID-19) has had devastating global health consequences and there is currently no cure and no licensed vaccine. Neutralizing antibodies (nAbs) to the causative agent of the disease, severe acute respiratory syndrome coronavirus-2 (SARS-CoV-2), represent potential prophylactic and therapeutic options and could help guide vaccine design. Indeed, a nAb to another respiratory virus, respiratory syncytial virus (RSV), is in widespread clinical use prophylactically to protect vulnerable infants [1]. Furthermore, nAbs prevent death from the emerging Ebola virus in macaques, even when given relatively late in infection, and thus have been proposed for use in humans in outbreaks [2,3]. Generally, nAbs with outstanding potency (“super-antibodies”) [4] can be isolated by deeply mining antibody responses of a sampling of infected donors. Outstanding potency together with engineering to extend antibody half-life from weeks to many months brings down the effective costs of Abs and suggests more opportunities for prophylactic intervention. At the same time, outstanding potency can permit anti-viral therapeutic efficacy that is not observed for less potent antibodies [4]. Here, we present the isolation of highly potent nAbs to SARS-CoV-2 and demonstrate their *in vivo* efficacy in a small animal model, suggesting their potential utility as a medical countermeasure.

To interrogate the antibody immune response against SARS-CoV-2 and discover nAbs, we adapted our pipeline to rapidly isolate and characterize monoclonal antibodies (mAbs) from convalescent donors (Fig. 1). Briefly, a cohort of previously swab-positive SARS-CoV-2 donors was recruited for peripheral blood mononuclear cell (PBMC) and plasma collection. In parallel, we developed both live replicating and pseudovirus neutralization assays using a HeLa-ACE2 (Angiotensin-Converting Enzyme-2) cell line that gave robust and reproducible virus titers. Convalescent serum responses were evaluated for neutralization activity against SARS-CoV-1 and SARS-CoV-2, and eight donors were selected for mAb discovery. Single antigen-specific memory B cells were sorted and their corresponding variable genes were recovered and cloned using a high-throughput expression system that enabled antibody expression and characterization in under two weeks. Promising mAbs were advanced for further biophysical characterization and *in vivo* testing.

**Figure 1.**
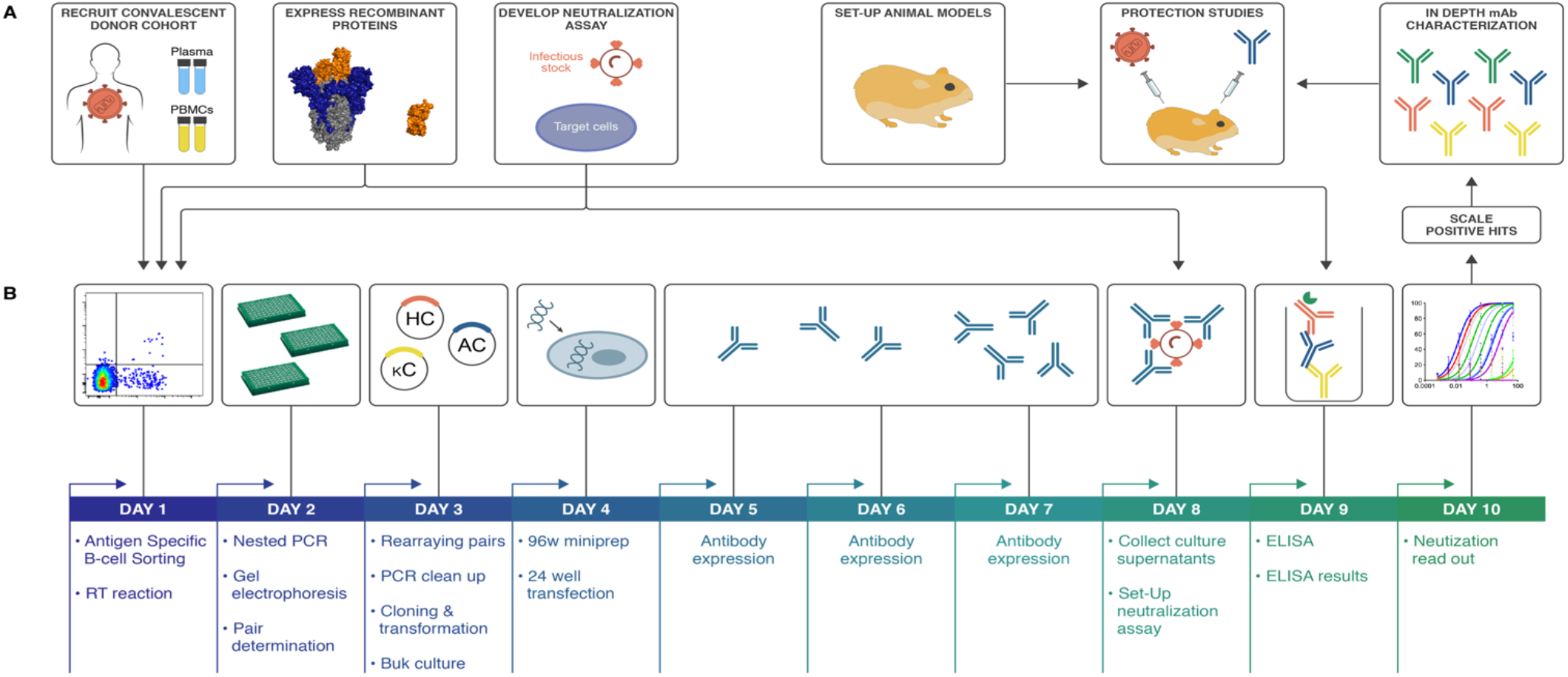
SARS-CoV-2 neutralizing antibody isolation strategy. (A) A natural infection cohort was established to collect plasma and PBMCs samples from individuals who recovered from COVID-19. In parallel, functional assays were developed to rapidly screen all plasma samples for SARS-CoV-2neutralizing activity. SARS-CoV-2 recombinant surface proteins were also produced to use as baits in single memory B-cell sorting and downstream functional characterization of isolated mAbs. Finally, a hamster animal model was set-up to evaluate mAb passive transfer protection. (B) The standard mAb isolation pipeline was optimized to allow high-throughput amplification, cloning, expression and functional screening of hundreds of unpurified Ab heavy and light chain pairs isolated from each of several selected neutralizers in only 10 days. Selected pairs were scaled-up to purify IgG for validation and characterization experiments. The most potent neutralizing mAb was selected to evaluate protection in the Syrian hamster model.

### Development of viral neutralization assays

Two platforms were established to evaluate plasma neutralization activity against SARS-CoV-2, one using replication-competent virus and another using pseudovirus (PSV). Vero-E6 cells were first used as target cells for neutralization assays, but this system gave poor virus titers for replicating virus (fig. S1A). To improve assay sensitivity, we established new target cells deriving from the HeLa cell line that stably expressed the cell surface ACE2 receptor. The HeLa-ACE2 target cell line gave reproducible titers and were used for the remainder of the study, with a comparison made between HeLa-ACE2 and Vero cells made in certain critical instances.

The live replicating virus assay used the Washington strain USA-WA1/2020 (BEI Resources NR-52281) and was optimized to a 384-well format to measure plaque formation. In parallel, a PSV assay was established for both SARS-CoV-1 and SARS-CoV-2 using murine leukemia virus (MLV)-based PSV [5]. The assay used single cycle infectious viral particles bearing firefly luciferase reporter for high-throughput screening. Unlike MLV-PSV, which buds at the plasma membrane, coronaviruses assemble in the ER-Golgi intermediate compartment, so the C-terminus of the SARS-CoV-1 Spike protein (S protein) contains an ER retrieval signal [6]. The alignment of SARS-CoV-1 and CoV-2 S proteins showed that this ER retrieval signal is conserved in SARS-CoV-2 (fig. S1B). To prepare high titers of infectious MLV-CoV-1 and SARS-CoV-2 PSV particles, various truncations of CoV-1 and CoV-2 S protein were carried out in which the ER retrieval signal was removed to improve exocytosis of the virus. Pseudoviron versions carrying SARS-CoV1-SΔ28 and SARS-CoV2-SΔ18S protein efficiently transduced ACE2-expressing target cells, but not control HeLa or A549 cells (fig. S1C). Control VSV-G pseudotyped virions showed a similar transduction efficiency in all target cells. Luciferase expression in transduced cells proved to be proportional to viral titer over a wide range (fig. S1D).

### Establishment of SARS-CoV-2 cohort

In parallel to the development of neutralization assays, a cohort was established in San Diego, California of 17 donors who had previously been infected with SARS-CoV-2 (Fig. 2A, fig. S2A, table S1). The cohort was 47% female and the average age was 50 years. Infection was determined as a positive SARS-CoV-2 PCR test from a nasopharyngeal swab. All donors also had symptoms consistent with COVID-19, and disease severity ranged from mild to severe, including intubation in one case, although all recovered. Donor plasma were tested for binding to recombinant SARS-CoV-2 and SARS-CoV-1 S and receptor binding domain (RBD) proteins, cell surface expressed spikes as well as for neutralization in both replicating virus and pseudovirus assays (Fig. 2B-D and fig. S2B). Binding titers to SARS-CoV-2 S protein varied considerably, reaching EC_50_s around 10_4_ with titers against the RBD about an order of magnitude less. Titers against SARS-CoV-1 S protein were notably less than for SARS-CoV-2 S protein and for SARS-CoV-1 RBD were only detected in a small number of donors. Neutralizing titers in the PSV assay varied over a wide range for SARS-CoV-2 (Fig. 2D) and were low or undetectable against SARS-CoV-1. Importantly, there was a strong linear relationship between RBD binding and pseudovirus neutralization (Fig. 2E). There was also a positive correlation between cell surface spike binding and live replicating virus neutralization (fig. S2C). The titers in the PSV assay were also largely reproduced in the replicating virus assay (fig. S2-3). In most later measurements, the PSV assay was preferred owing to its higher reproducibility and throughput. Based on the plasma neutralization results, we prioritized donors CC6, CC12 and CC25 for antigen-specific B cell sorting from cryopreserved PBMCs.

**Figure 2.**
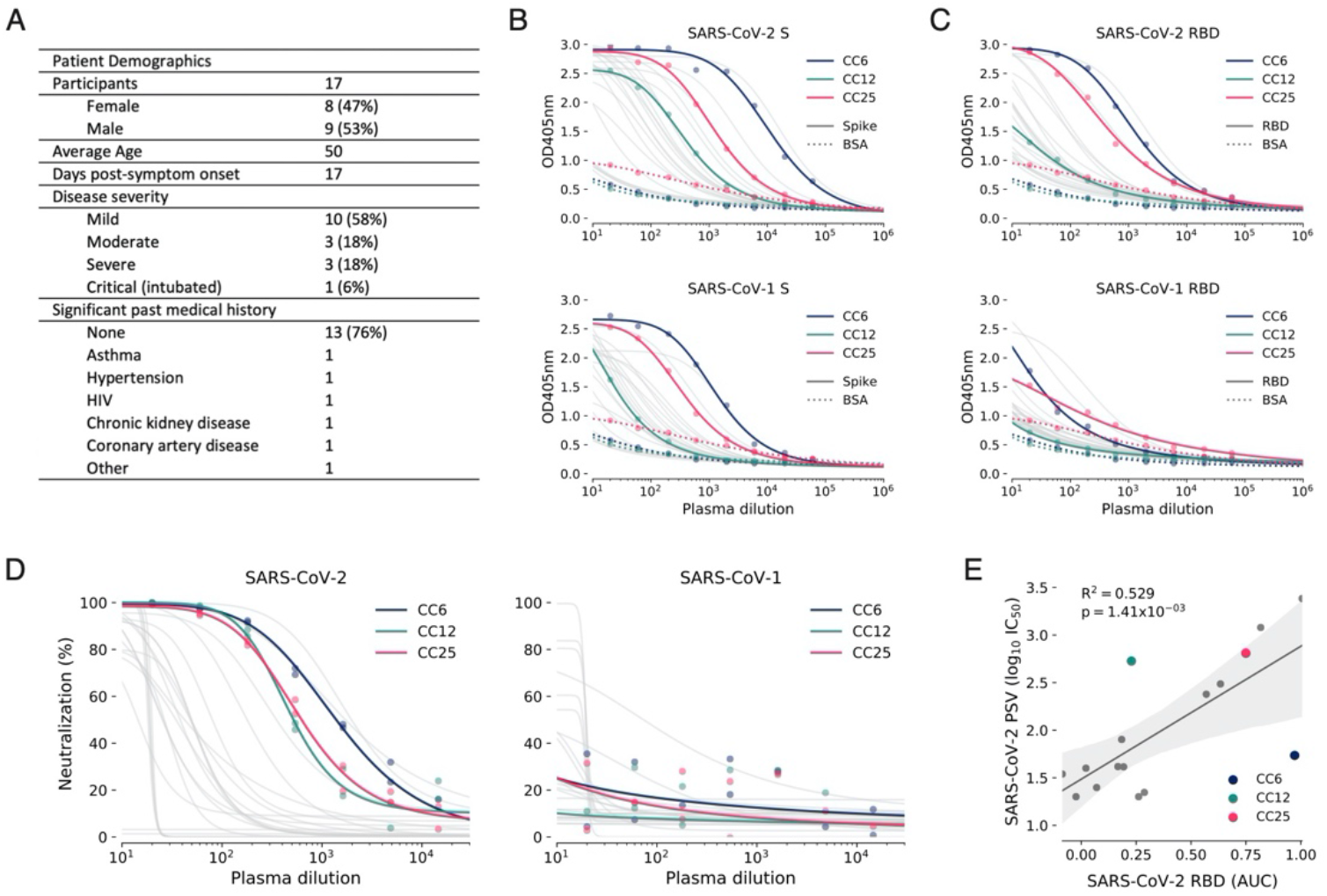
COVID-19 cohort functional screening. (**A**) Demographics of the UCSD COVID-19 cohort (CC) participants. CC plasma were tested for binding to SARS-CoV-1 and SARS-CoV-2 S protein (**B**) and RBD subunit (**C**) by ELISA. Background binding of plasma to BSA-coated plates is represented by a dashed line. (**D**) Plasma were also tested for neutralization of pseudotyped (PSV) SARS-CoV-1 and SARS-CoV-2 virions. (**E**) Correlation between PSV SARS-CoV-2 neutralization and RBD subunit ELISA binding area-under-the-curve (AUC). AUC was computed using Simpson’s rule. The 95% confidence interval of the regression line is shown in light grey and was estimated by performing 1,000 bootstrap re-samplings. R2 and p values of the regression are also indicated. CC participants from whom mAbs were isolated are specifically highlighted in dark blue (CC6), pine green (CC12) and hot pink (CC25).

### Antibody isolation and preliminary functional screens for downselection

PBMCs from donors CC6, CC12, and CC25 were stained for memory B cells markers (CD19+/IgG+) and both Avi-tag biotinylated RBD and SARS-CoV-2 S antigen baits before single-cell sorting. S+ and S+/RBD+ memory B cells were present at an average frequency of 3.2% and 0.7%, respectively, across the three donors (fig.S4A). In total, 1721 Ag+ memory B cells were sorted to rescue native heavy and light chain pairs for mAb production and validation (fig. S4B).

Across the 3 donors, a total of 1043 antibodies were cloned and expressed, which represents a 65% PCR recovery of paired variable genes and >86% recovery of fully functional cloned genes (fig. S4C). The bulk-transformed ligation products for both the heavy chain and light chain were transfected and tested for binding to RBD and S protein, and for neutralization in the SARS CoV-2 pseudovirus assay using HeLa-ACE2 target cells (Fig. 3 and fig. S5).

**Figure 3.**
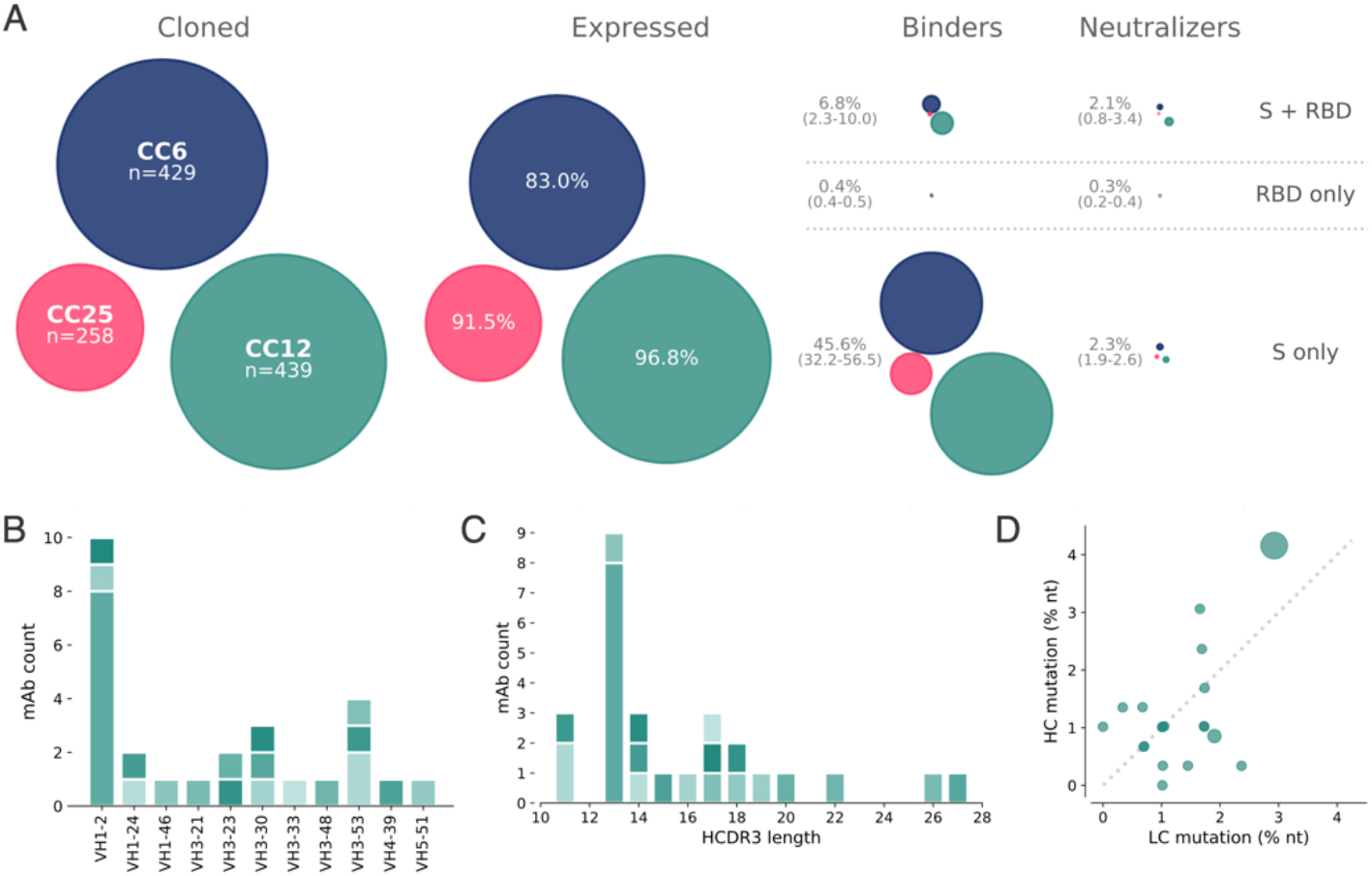
Antibody isolation and functional screening for SARS-CoV neutralization and antigen binding. (**A**) Antibody downselection process from 3 donors, presented as bubble plots. The areas of the bubbles for each donor are sized based on the number of antibodies that were cloned and transfected, then scaled according to the number that were positive in subsequent assays. All antibodies that expressed at measurable levels were tested for binding to S protein and RBD to determine their specificity, and then screened for neutralization. (**B**) VH gene distribution of downselected mAbs. Antibodies are colored by their respective clonal lineages. (**C**) Heavy chain CDR3 lengths of downselected mAbs. Antibodies are colored by their respective clonal lineages. (**D**) Mutation frequency of downselected mAb lineages. Bubble position represents the mean mutation frequency for each lineage, with bubble area proportional to the lineage size.

On average, 92% of the transfected pairs resulted in IgG expression. Of these, 46% showed binding only to S protein while 6.8% bound to both S and RBD proteins and 0.2% bound only to RBD. The supernatants were also screened for binding to an unrelated HIV antigen (BG505 SOSIP) to eliminate non-specific or polyreactive supernatants. The supernatants were next evaluated for neutralization activity using SARS-CoV-2 and SARS-CoV-1 pseudoviruses. Strikingly, a small proportion of the binding antibodies showed neutralization activity and that activity was equally distributed between RBD+/S+ and S+ only binders despite a much larger number of S+ only binding supernatants (Fig. 3A). These data indicate that viral infection generates a strong response against the non-RBD regions of S protein, but only a small proportion of that response is neutralizing. In contrast, there are fewer RBD binding antibodies but a larger proportion of these are capable of neutralizing SARS-CoV-2 pseudovirus. Antibodies that tested positive for neutralization in the high-throughput screening were sequence confirmed and advanced for expression at large scale for additional characterization.

Sequencing of these Abs identified 19 distinct lineages, with 17 containing a single member (table S2). VH1 and VH3-gene families were notably prominent in these Abs and there was a diversity of CDR3 lengths (Fig. 3B-C). There was one prominent example of a clonally expanded lineage, with 8 recovered clonal members that averaged 4.3% and 2.8% mutations from germline at the nucleotide level in the heavy chain and light chain, respectively (Fig. 3D). The remaining clones were relatively unmutated, averaging just above 1% mutation at the nucleotide level suggesting that these antibodies were primed by the ongoing COVID infection and likely not recalled from a previous endemic human coronavirus (HCoV) exposure.

All antibodies that were expressed at scale were evaluated in a standard ELISA-based polyreactivity assay with solubilized CHO membrane preparations, ssDNA and insulin, and none were found to be polyreactive (fig. S6).

### Functional activity of downselected antibodies

The antibody hits that were identified in the high-throughput screening were next evaluated for epitope specificity by bio-layer interferometry (BLI) using S and RBD proteins as capture antigens. The antigens were captured on anti-HIS biosensors before addition of a saturating concentration (100 μg/mL) of antibodies that were then followed by competing antibodies at a lower concentration (25 μg/mL). Accordingly, only antibodies that bind to a non-competing site would be detected in the assay. Among the antibodies evaluated, the results reveal 3 epitope bins for RBD (designated as RBD-A, RBD-B, and RBD-C) and 3 epitope bins for the S protein (designated as S-A, S-B, and S-C) (Fig. 4A and fig. S7). Interestingly, the mAb CC12.19 appears to compete with antibodies targeting two different epitopes, RBD-B and S-A, which might indicate that this mAb targets an epitope spanning RBD-B and S-A. To evaluate epitope specificities further, we next assessed binding of the antibodies to extended RBD-constructs with subdomains (SD) 1 and 2, including RBD-SD1 and RBD-SD1-2, and the N-terminal domain (NTD) (Fig. 4B and fig.S8A-B). None of the antibodies showed binding to the NTD. CC12.19 binds to all the other constructs, which supports the epitope binning data described in Figure 4A. The other antibodies grouped in the S-A epitope bin that compete with CC12.19 show either no binding to RBD or RBD-SD constructs (CC12.20 and CC12.21) or do show binding to RBD-SD1 and RBD-SD1-2 but not RBD (CC12.23). These data suggest two competing epitopes within the S-A epitope bin; one that is confined only to the S protein and the other that includes some element of RBD-SD1-2. This interpretation will require further investigation by structural studies.

**Figure 4.**
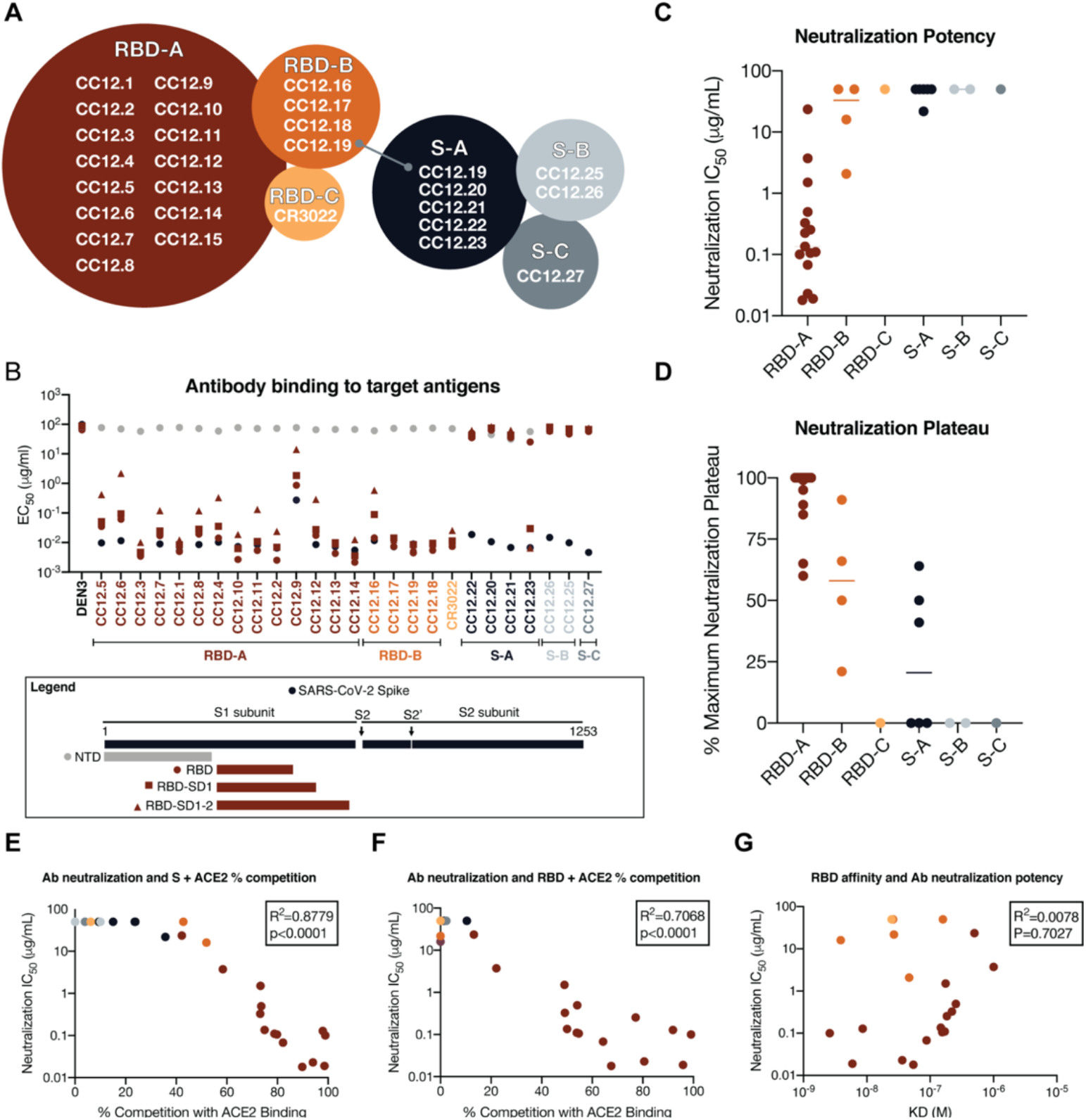
Antibody functional activity by epitope specificities. Monoclonal antibody epitope binning was completed using RBD and SARS-CoV-2 S protein as target antigens. (**A**) A total of three non-competing epitopes for RBD (RBD-A, RBD-B, and RBD-C) and three non-competing epitopes for S (S-A, S-B, and S-C) were identified. (**B**) MAbs were evaluated for binding to different target antigens (S, N-terminal domain (NTD), RBD, RBD-SD1, and RBD-SD1-2) by ELISA and apparent EC_50_s are reported in *μ*g/ml. (**C**) MAbs were evaluated for neutralization on SARS-CoV-2 pseudovirus and HeLa-ACE2 target cells. Antibodies are grouped according to epitope specificities and neutralization IC_50_ values are reported in *μ*g/ml. (**D**) The maximum plateaus of neutralization (MPN) are reported for each mAb and grouped by epitope specificity. MAbs were mixed with (**E**) RBD or (**F**) S protein and measured for binding to HeLa-ACE2 target cells as a measure of competition to the cell surface ACE-2 receptor. (**G**) Monoclonal antibody neutralization potencies (IC_50_, *μ*g/ml) are plotted compared to dissociation constants (KD, M) measured by surface plasmon resonance (SPR) to RBD target antigen.

We next evaluated the mAbs for neutralization activity against SARS-CoV-2 pseudovirus. The neutralization IC_50_ potencies of these antibodies are shown in Fig. 4C and their associated maximum plateaus of neutralization (MPNs) are shown in Fig. 4D. A comparison of neutralization potency between pseudovirus (fig. S8C) and live replicating virus (fig. S8D) are also included. Notably, the most potent neutralizing antibodies were those directed to RBD-A epitope. In comparison, antibodies directed to RBD-B have markedly higher IC_50_ values and also plateau below 100%. The antibodies that do not bind to RBD and are directed to epitopes on S protein all show poor neutralization potencies and MPNs well below 100%.

To evaluate whether the RBD-A epitope might span the ACE2 binding site, we next performed cell surface competition experiments. Briefly, antibodies were premixed with biotinylated S (Fig. 4E) or RBD (Fig. 4F) proteins at a molar ratio of 4:1 of antibodies to target antigen. The mixture was then incubated with the HeLa-ACE2 cell line and the percent competition against ACE2 receptor was recorded by comparing percent binding of the target antigen with and without antibody present (fig. S8E). The data indicate that the antibodies targeting the RBD-A epitope compete best against the ACE2 receptor and that the neutralization IC_50_ correlates well with the percent competition for ACE2 receptor binding for both S protein (Fig. 4E) and for RBD (Fig. 4F). We also assessed the affinity of all RBD-specific antibodies to soluble RBD by surface plasmon resonance (SPR) and found a poor correlation between affinity and neutralization potency (Fig. 4G and fig. S9), however, the correlation is higher when limited to antibodies targeting the RBD-A epitope. The lack of a correlation between RBD binding and neutralization for mAbs contrasts with the strong correlation described earlier for serum RBD binding and neutralization. Overall, the data highlight epitope RBD-A as the preferred target for eliciting neutralizing antibodies and that corresponding increases in affinity of mAbs to RBD-A will likely result in corresponding increases in neutralization potency. The data also suggest that less potent nAbs targeting RBD-B that compete with RBD-A nAbs could, at least in principle, pose challenges for the elicitation of RBD-A nAbs.

### Passive transfer of neutralizing antibodies and SARS-CoV-2 challenge in Syrian hamsters

To investigate the relationship between *in vitro* neutralization and protection *in vivo* against SARS-CoV-2, we selected two mAbs for passive transfer/challenge experiments in a Syrian hamster animal model based on a summary of the nAb data (table S3 and fig.S10). The experimental design for the passive transfer study is shown in Fig. 5A. In the first experiment, we tested nAb CC12.1, which targets the RBD-A epitope and has an in vitro IC_50_ neutralization of 0.019 μg/mL against pseudovirus and in the second we tested nAb C12.23, which targets the S-B epitope with an IC_50_ neutralization of 22 μg/mL. In both experiments an unrelated antibody to dengue virus, Den3, was used as a control. The anti-SARS-CoV-2 nAbs were delivered at 5 different concentrations to evaluate dose-dependent protection starting at 2 mg/animal (average of 16.5 mg/kg) at the highest dose and 8 μg/animal at the lowest dose. The Den3 control antibody was delivered at a single dose of 2 mg/animal. Sera were collected from each animal 12 hours post IP infusion of the antibody and all animals subsequently were challenged with a dose of 1×10_6_ PFU of SARS-CoV-2 (USA-WA1/2020) by intranasal administration 12h post antibody infusion (Fig. 5A).

**Figure 5.**
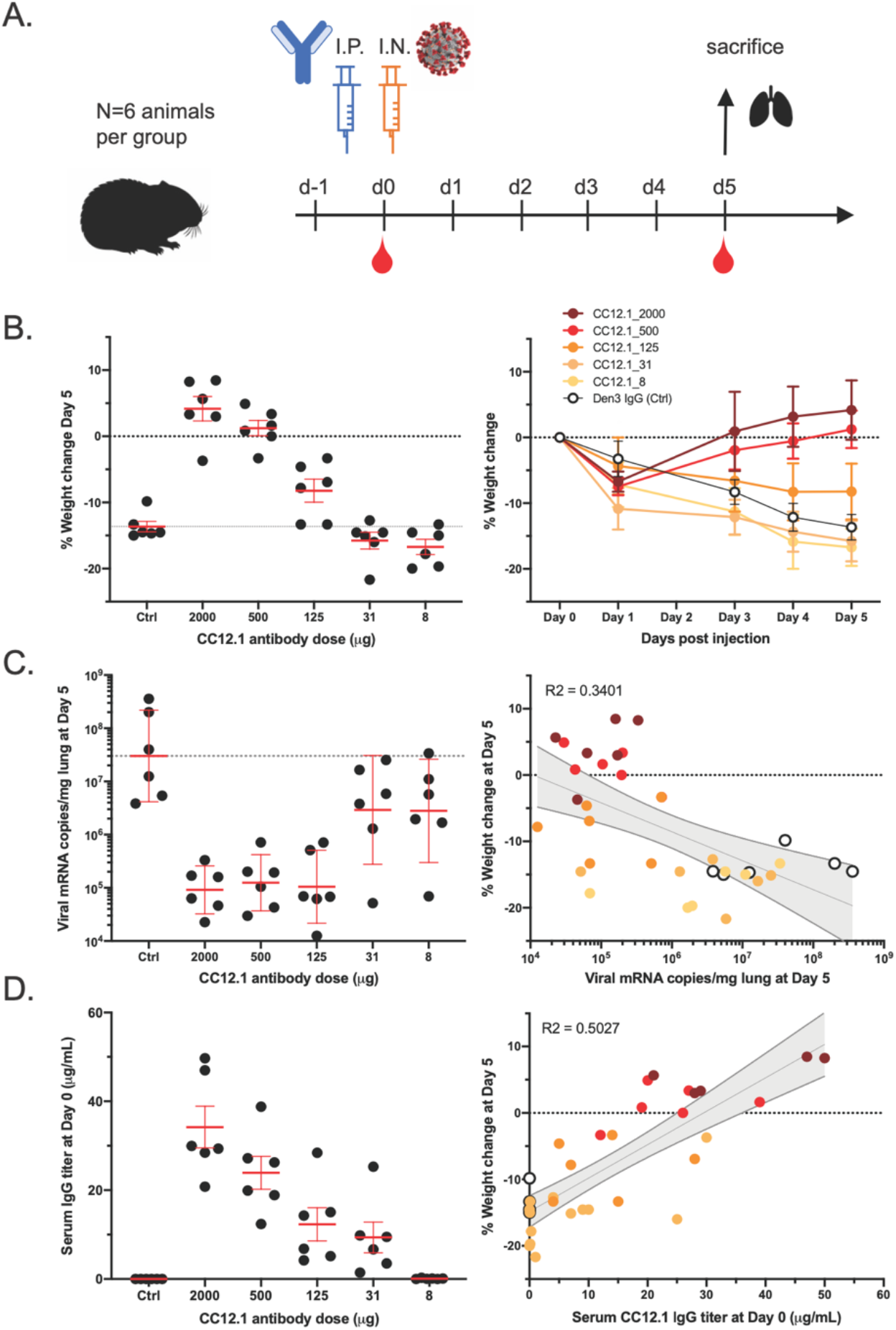
A potent SARS-CoV-2 RBD-specific neutralizing mAb protects against disease progression and lung viral burden in Syrian hamsters. (**A**) SARS-CoV-2-specific human neutralizing mAb CC12.1 isolated from natural infection was injected intraperitoneally into Syrian hamsters at a starting dose of 2 mg/animal (on average 16.5 mg/kg) and subsequent serial 4-fold dilutions. Control animals received 2 mg of a dengue-specific human IgG1 (Den3). Each group of 6 animals were challenged intranasally 12h post-infusion with 1X106 PFU of SARS-CoV-2. Serum was collected at the time of challenge (Day 0) and Day 5, and their weight monitored as an indicator of disease progression. On day 5, lung tissue was collected for viral burden assessment. (**B**) Percentage weight change was calculated from day 0 for all animals at all time points. (**C**) Viral load as assessed by Q-PCR from lung tissue at day 5. (**D**) Serum titers of the passively administered mAb, as assessed by ELISA at the time of challenge (12h after i.p administration). Correlation analyses with 95% confidence intervals indicated in grey shade. R^2^ values are also indicated.

Syrian hamsters typically clear virus within one week after SARS-CoV-2 infection [7]. Accordingly, the hamsters were weighed as a measure of disease due to infection. Lung tissues were collected to measure viral load on day 5. A data summary is presented in Fig. 5B and fig. S11A for animals that received CC12.1, which targets the RBD-A epitope. The control animals that received Den3 lost on average 13.6% of body weight at 5 days post virus challenge. In comparison, the animals that received the neutralizing RBD-A antibody at a dose of 2 mg (average of 16.5 mg/kg) or 500 μg (average of 4.2 mg/kg) exhibited no weight loss. However, animals that received a dose of 125 μg (average of 0.9 mg/kg) had aon average 8% loss of body weight, while animals that received a dose of 31 μg/mL (0.2 mg/kg) and 8 μg/mL (0.06 mg/kg) lost 15.8% and 16.7% of body weight, respectively. We note these animals showed a trend for greater weight loss than control animals but this did not achieve statistical significance (table S4). Given concerns about antibody-mediated enhanced disease in SARS-CoV-2 infection, this observation merits further attention using larger animal group sizes. The weight loss data are further corroborated by quantification of lung viral load measured by real-time PCR (Fig. 5C) and showed a moderate correlation to weight loss. The data indicate comparable viral loads between the three higher doses (2 mg, 500 μg, and 125 μg) of nAbs. In contrast, equivalent viral loads were observed between the control group receiving Den3 and the low dose groups receiving 31 μg and 8 μg of nAb. In contrast to the nAb to RBD-A, the less potent and incompletely neutralizing antibody to the S-B epitope showed no evidence of protection at any concentration compared to the control animals (fig. S11B).

To determine the antibody serum concentrations that may be required for protection against SARS-CoV-2 *in vivo*, we also measured the antibody serum concentrations just prior to intranasal virus challenge (Fig. 5D). The data highlight that an antibody serum concentration of approximately 22 μg/mL of nAb (1160 × PSV neutralization IC_50_) enables full protection and a serum concentration of 12 μg/mL (630 × PSV neutralization IC_50_) is adequate for 50% reduced disease as measured by weight loss. The effective antibody concentration required at the site of infection to protect from disease remains to be determined. Sterilizing immunity at serum concentrations that represent a large multiplier of the *in vitro* neutralizing IC_50_ is observed for many viruses [8].

## Discussion

Using a high-throughput rapid system for antibody discovery, we isolated more than 1,000 mAbs from 3 convalescent donors by memory B cell selection using SARS-CoV-2 S or RBD recombinant proteins. About half of the mAbs isolated could be expressed and also bind effectively to either S and/or RBD proteins. The overwhelming majority of these mAbs were not functional in either pseudovirus or replicating virus neutralization assays, which were largely internally consistent with one another. The low frequency of nAbs highlights the value of deep mining of responses to access the most potent Abs [4].

A range of nAbs were isolated to different sites on the S protein. The most potent Abs, reaching low double digit ng/mL IC_50_ values in PSV assays, are targeted to a site that, judged by competition studies, overlaps with the ACE2 binding site. These Abs show no neutralization of SARS-CoV-1 PSV, as may be anticipated given the differences in ACE2 contact residues between the two viruses (fig. S12). Abs directed to the RBD but not competitive with soluble ACE2, (although they may be competitive in terms of an array of membrane-bound ACE2 molecules interacting with an array of spike proteins on a virion), are notably less potent neutralizers and tend to show incomplete neutralization, plateauing at around or less than 50% neutralization. Similar behavior is observed for Abs to the S protein that are not reactive with recombinant RBD. The cause(s) of these incomplete neutralization phenomena is unclear but presumably originates in some spike protein heterogeneity, either glycan-or conformationally based. In any case, the RBD-A nAbs that directly compete with ACE2 are clearly the most preferred for prophylactic and therapeutic applications, and as reagents to define nAb epitopes for vaccine design.

In terms of nAbs as passive reagents, the observation of efficacy of a potent anti-RBD nAb in vivo in Syrian hamsters is promising in view of the positive attributes of this animal model [9] and suggests that human studies are merited. The failure of the non-RBD S-protein nAb to protect in the animal model is consistent with its lower potency and, likely most importantly, its inability to fully neutralize challenge virus. In the context of human studies, improved potency of protective nAbs by enhancing binding affinity to the RBD epitope identified, improved half-life and reduced Fc receptor binding to minimize potential antibody dependent enhancement (ADE) effects, should they be identified as concerning, are all antibody engineering goals to be considered. If ADE is found for SARS-CoV-2 and operates at sub-neutralizing concentrations of neutralizing antibodies as it can for dengue virus [10] then it will be important to carefully define the full range of nAb epitopes on the S protein as we have begun here.

The nAbs described have remarkably little SHM, typically one or two mutations in the VH gene and one or two in the VL gene. Such low SHM may be associated with the isolation of the nAbs relatively soon after infection, and perhaps before affinity-maturation has progressed. Low SHM has also been described for potent nAbs to Ebola virus, respiratory syncytial virus (RSV), Middle East Respiratory Syndrome coronavirus (MERS-CoV) and yellow fever virus [11–14] and may indicate that the human naïve repertoire is often sufficiently diverse to respond effectively to many pathogens with little mutation. Of course, nAb efficacy and titer may increase over time as described for other viruses and it will be interesting to see if even more potent nAbs to SARS-CoV-2 evolve in our donors in the future.

What do our results suggest for SARS-CoV-2 vaccine design? In the first instance, the results suggest a focus on the RBD and indeed strong nAb responses have been described by immunizing mice with a multivalent presentation of RBD [15]. The strong preponderance of non-neutralizing antibodies and very few nAbs to S protein that we isolated could arise for a number of reasons including: (1) the recombinant S protein that we used to select B cells is a poor representation of the native spike on virions. In other words, there may be many nAbs to S but we failed to isolate them because of the selecting antigen, (2) the recombinant S protein that we used is close to native but non-neutralizing antibodies bind to sites on S that do not interfere with viral entry, (3) the S protein in natural infection disassembles readily generating a strong Ab response to “viral debris” that is non-neutralizing because the antibodies recognize protein surfaces that are not exposed on the native spike. Importantly, the availability of both neutralizing and non-neutralizing antibodies generated in this study will facilitate evaluation of S protein immunogens for presentation of neutralizing and non-neutralizing epitopes and promote effective vaccine design.

In summary, we describe the very rapid generation of neutralizing antibodies to a newly emerged pathogen. The antibodies can find clinical application and will aid in vaccine design.

## Acknowledgements

We thank Tom Gilman, Andrea Salazar and Biosero for their contribution to high-throughput pipeline generation. We also thank Barney Graham, Jason S. McLellan and Ian A. Wilson for reagents. We thank Laura Walker for valuable manuscript discussions.

This work was supported by the NIH CHAVD (UM1 AI44462 to B.B., D.S. and D.R.B.) awards, the IAVI Neutralizing Antibody Center, the Bill and Melinda Gates Foundation (OPP 1170236 to D.R.B.) and (OPP 1206647 to D.R.B and R.A.), and (OPP1196345/ INV-008813 to DS and DRB).

This work was also supported by the John and Mary Tu Foundation and the Pendleton Foundation.

