## Supplementary Material for "Rapid isolation of potent SARS-CoV-2 neutralizing antibodies and protection in a small animal model"

Materials and Methods
Tables S1 – S2
Figures S1 – S12
Supplementary References (16 – 25)

#### Materials and Methods

HeLa-hACE2 Stable Cell Line

HeLa-hACE2 and A549-hACE2 cells were generated through transduction of human ACE2 lentivirus. pBOB-hACE2 construct was co-transfected into HEK293T cells along with lentiviral packaging plasmids pMDL, pREV, and pVSV-G (Addgene) by Lipofectamine 2000 (Thermo Fisher Scientific, 11668019) according to the manufacturer's instructions. Supernatants were collected 48 h after transfection, then were transduced to pre-seeded HeLa or A549 cells. 12 h after transduction, stable cell lines were collected, scaled up and stored for neutralization assay.

Virus generation

Vero E6 cells (ATCC CRL-1586) were plated in a T225 flask with complete DMEM (Corning 15-013-CV) containing 10% FBS, 1X PenStrep (Corning 20-002-CL), 2 mM L-Glutamine (Corning 25-005-CL) overnight at 37C 5% CO2. The media in the flask was removed and 2 mL of SARS-CoV-2 strain USA-WA1/2020 (BEI Resources NR-52281) in complete DMEM was added to the flask at an MOI of 0.5 and was allowed to incubate for 30 minutes at 34˚C 5% CO2. After incubation, 30 mL of complete DMEM was added to the flask. The flask was then placed in a 34°C incubator at 5% CO2 for 5 days. On day 5 post infection, the supernatant was harvested and centrifuged at 1,000× g for 5 minutes. The supernatant was filtered through a 0.22 μM filter and stored at -80˚C.

SARS-CoV-2 focus reduction neutralization test (FRNT)

HeLa-ACE2 cells were plated in 12 μL complete DMEM at a density of 2×103 cells per well. In a dilution plate, plasma or mAb was diluted in series with a final volume of 12.5 μL. 12.5 μL of SARS-CoV-2 was added to the dilution plate at a concentration of 1.2E4 pfu/mL. The dilution plate was covered and incubated at 34°C 5% CO2 for 1h. After incubation, the media remaining on the 384-well cell plate was removed and 25μL of the virus plasma/mAb mixture was added to the 384-well cell plate. The plate was incubated for 20hat 34°C 5% CO2. After incubation, the plate was fixed with 25 μL of 8% Formaldehyde for 1h at 34°C 5% CO2. The plate was then washed three times with 100 μL of 1×XPBS 0.05% tween. 12.5 μL of human polyclonal sera diluted 1:500 in Perm/Wash buffer (BD Biosciences 554723) were added to the plate and incubated at RT for 2h. The plate was washed three times with 100 μL of 1×XPBS 0.05% tween. 12.5 μL of Peroxidase goat anti-human Fab (Jackson Scientific) were diluted 1:200 in Perm/Wash buffer then added to the plate and incubated at RT for 2h. The plate was then washed three times with 50 μL of 1×X PBS 0.05% tween. 12.5 μL of Perm/Wash buffer was added to the plate then incubated at RT for 5 min. The Perm/Wash buffer was removed and TrueBlue peroxidase substrate was immediately added (Sera Care 5510-0030). Infected cell non-linear regression curves were analyzed using Prism 8 software to calculate $EIC_{50}$ values.

Pseudovirus (PSV) Assay

MLV-gag/pol and MLV-CMV-Luciferase plasmids were co-transfected with full-length or truncated SARS-CoV-2 and SARS-CoV-2 plasmid, respectively, with transfection reagent Lipotransfectmine 2000 in HEK293T cells. After 48 h of transfection, supernatants containing pseudotyped virus were collected and frozen at -80°C for long-term storage. Serially diluted plasma or mAbs were incubated with pseudovirus at 37°C for 1 h, then transferred onto HeLa-hACE2 cells in 96-well plates at 10,000 cells/well (Corning, 3688). After 48 h of incubation, supernatants were removed, HeLa-hACE2 cells were then lysed in luciferase lysis buffer (25mM Gly-Gly pH 7.8, 15mM MgSO4, 4 mM EGTA, 1% Triton X-100). Luciferase activity was measured by adding Bright-Glo (Promega, PR-E2620) according to the manufacturer's instructions. Plasma or mAbs were tested in duplicate wells. Neutralization ID50 or $IC_{50}$

titers were calculated using “One-Site Fit LogIC$_{50}$” regression in GraphPad Prism 8.0.

Cohort information

De-identified PBMC and plasma were kindly provided through the “Collection of Biospecimens from Persons Under Investigation for 2019-Novel Coronavirus Infection to Understand Viral Shedding and Immune Response Study” UCSD IRB# 200236. Protocol was approved by the UCSD Human Research Protection Program.

Whole Virus ELISA

High binding plates (Corning 3700) were coated with 12.5 µL of Galanthus Nivalis Lectin (GNL; Vector Laboratories L-1240-5) at 10 µg/mL and incubated overnight at 4˚C. The GNL was removed and 12.5 µL of SARS-CoV-2 was added to the plate at a concentration of 2E6 pfu/mL then incubated for 24 hours at 4˚C. 12.5 µl of 8% Formaldehyde was added to a final concentration of 4% then incubated at RT for 1 hour. The plate was then washed three times with 100 µL of 1x PBS supplemented with 0.05% tween. 50 µL of 3% BSA were added to the plate and incubated at RT for 2 hours. The BSA was removed and 12.5 µL of plasma or mAb diluted in series was added to the plate then incubated at RT for 1.5 hours. The plate was then washed three times with 100 µL of 1x PBS supplemented with 0.05% tween. 12.5 µL of alkaline phosphatase conjugated goat anti-human Fc antibody (Jackson Immunoresearch 109-055-098) diluted 1:2000 was added to the plate and incubated for 1 hour at RT. The plate was then washed three times with 100 µL of 1x PBS supplemented with 0.05% tween. 12.5 µL of phosphatase substrate (SIGMA-ALDRICH S0942) were added to the plate. Non-linear regression curves were analyzed using Prism 8 software to calculate EC50 values.

Plasmid construction for full-length and recombinant soluble proteins

To generate full-length SARS-CoV-1 (1255 amino acids; GenBank: AAP13567) and SARS-CoV-2 (1273 amino acids; GenBank: MN908947) spike plasmids, genes were synthesized from GeneArt (Life Technologies), cloned into mammalian expression vector phCMV3 (Genlantis, USA) using PstI and BamH restriction sites. Soluble S ectodomain protein SARS-CoV-1 (residue 1-1190) and SARS-CoV-2 (residue 1-1208) encoding plasmids were constructed by PCR amplifications and Gibson assembly cloning into vector phCMV3. To stabilize soluble S proteins in prefusion state and to improve trimerization, double proline substitutions in the S2 subunit, replacement of a furin cleavage site in SARS-CoV-2 (residues 682–685), and S2 cleavage site in SARS-CoV-1 (residues 664–667) with “GSAS” and incorporation of a C-terminal T4 fibritin trimerization motif, as described previously [16,17]. An HRV-3C protease cleavage site, a 6X HisTag, and an AviTag spaced by GS-likers were added to the C-terminus to aid purification strategies. To generate gene fragments encoding SARS-CoV-2 N-terminal domain-NTD (residue 1-290), receptor-binding domain-RBD (residue 332-527), RBD-SD1 (residue 320-591), and RBD-SD1-2 (residue 320-681) subdomains, PCR-amplifications were carried out from the SARS-CoV-2 plasmid and gene fragments were cloned in frame with the original secretion signal or the Tissue Plasminogen Activator (TPA) leader sequence. A similar design strategy was used to construct the SARS-CoV-1-RBD (residue 319-513) gene encoding plasmid.

Flow cytometry based cell surface SARS-CoV-1/CoV-2 spike binding assay

Binding of mAbs/sera to the HEK293T cell-surface expressed SARS-CoV-1 and SARS-CoV-2 spikes was performed as described previously [18]. Briefly, HEK293T cells were transfected with plasmids encoding full-length SARS-CoV-1 or SARS-CoV-2 spikes and incubated for 36-48 hours at 37 °C. Post incubation cells were trypsinization to prepare single cell suspension and were distributed into 96-well plates. 50ul/well of 3-fold serial titrations of mAbs starting at 10 µg/mL or serum samples starting at 1:30 dilution were added to transfected cells. The Abs were incubated with cells for 1 hr on ice. The plates were washed twice in FACS buffer (1X PBS, 2% FBS, 1mM EDTA) and stained with 50ul/well of 1:200 dilution of R-phycoerythrin (PE)-conjugated mouse anti-human IgG Fc antibody (SouthernBiotech) and 1:1000 dilution of Zombie-NIR viability dye (BioLegend). After another two washes, stained cells were analyzed using flow cytometry (BD Lyrics cytometers), and the binding data were generated by calculating the percent (%) PE-positive cells for antigen binding using FlowJo 10 software. CR3022, a SARS-CoV-2 spike binding antibody, and unrelated antibody, DEN3, were respectively positive and negative controls for the assay.

Protein expression and purification

To express the soluble S ectodomain protein SARS-CoV-1, SARS-CoV-2 and their truncated protein versions, protein-encoding plasmids were transfected into FreeStyle293F cells (Thermo Fisher) at a density of approximately 1 million cells/mL. For large scale production, we mixed 350ug plasmids with 16 mL transfectagro™ (Corning) in a conical tube and filtered with 0.22um Steriflip™ Sterile Disposable Vacuum Filter Units (MilliporeSigma™). In another conical tube, we added 1.8mL 40K PEI (1 mg/mL) into 16 mL transfectagro™ and mixed briefly. The premixed 40K PEI- transfectagro™ solution was gently poured into the filtered plasmid solution. The solution was thoroughly mixed by inverting the tube several times. The mixture rested at room temperature for 30 mins and was poured into 1L FreeStyle293F cell culture. After 5 days, the cells were removed from the supernatant by centrifuging at 3500 rpm for 15mins. The supernatant was filtered in a glass bottle with the 0.22um membrane and kept in 4-degree storage before loading into the columns. The His-tagged proteins were purified with the HisPur Ni-NTA Resin (Thermo Fisher). To eliminate the nonspecific binding proteins, each column was washed with at least 3 bed volumes of wash buffer (25 mM Imidazole, pH 7.4). To elute the purified proteins from the column, we loaded 25mL of the elution buffer (250mM Imidazole, pH 7.4) at slow gravity speed (~4 sec/drop). By using Amicon tubes, we buffer exchanged the solution with PBS and concentrated the proteins. The proteins were further purified by size-exclusion chromatography using Superdex 200 (GE Healthcare). The selected fractions were pooled and concentrated again for the further use.

Recombinant Protein ELISAs

6x-His tag monoclonal antibody (Invitrogen, UA280087) was coated onto high-binding 96-well plates (Corning, 3690) at 2 µg/mL overnight at 4 °C. After washing, plates were blocked with 3% BSA in PBS for 1h. Then his-tag recombinant RBD and S protein were captured at 1µg/mL in 1% BSA and incubated for 1h at RT. After washing, serially diluted mAbs were added into wells and incubated for 1h at RT. Detection was measured with alkaline phosphatase-conjugated goat anti-human IgG Fcg (Jackson ImmunoResearch) at 1:1000 dilution for 1h. After the final wash, phosphatase substrate (Sigma-Aldrich) was added into wells. Absorption was measured at 405 nm.

12.5 ul of 6x-His tag monoclonal antibody (Invitrogen UA280087) were coated onto high binding plates at 2µg/mL overnight at 4°C. The plate was washed three times with 100 ul of 1xPBS/0.05% Tween. The recombinant RBD and S protein derived from SARS-CoV-1 and SARS-CoV-2 were diluted with 1xPBS/1%BSA to final concentrations of 2 µg/mL or 5 µg/mL and then coated onto the plate and incubated at RT for 2h. The plate was washed three times with 100 ul of 1xXPBS/0.05% Tween and then blocked with 100 ul of 3% BSA at RT for 2 hours and then the BSA was removed. Either serially diluted plasma samples or isolated mAbs were added onto the plate and incubated for 1.5 hours at RT. Wells were then incubated with secondary anti-human IgG Fcγ antibody diluted at 1:2000 and incubated at RT for 1 hour. The plate was then washed three times with 100 µL of 1xPBS/0.05% tween. 12.5uL of phosphatase substrate (SIGMA-ALDRICH S0942) were added to the plate and optical density (OD) was measured at 405 nm. Plasma or mAbs were tested in duplicate or triplicate wells. Non-linear regression curves were analyzed using Prism 8 software to calculate $IC_{50}$ values.

Isolation of SARS-2 S-specific mAbs

Sorting of antigen-specific memory B cells was performed as previously described [19,20]. The process was adapted for high-throughput such that each step could be performed in a 96-well format. Fluorescent-labeled antibodies recognizing cell surface markers were purchased from BD Biosciences. AVI-tagged SARS-2 S and RBD proteins were produced, purified, labeled with biotin (Avidity), and coupled to streptavidin-AF647, streptavidin-AF488 (Thermo Fisher), and streptavidin-BV421 (BD Biosciences), as previously described [20] at 2:1 and 4:1 molecular ratio respectively 30 min prior to staining. Cells were first labeled with antibodies for surface markers together with probes (200 nM final) for 30min in sort buffer (PBS 1% FBS, 2.5M EDTA, 25mM Hepes) on ice. Cells were then stained with the Live/Dead Fixable Aqua Dead Cell Stain (Thermo Fisher) for 15 minutes on ice according to the manufacturer's instructions. Single antigen-specific (S+ and RBD+) memory B cells (CD3-CD4-CD8-CD14-CD19+IgD-IgG+) were sorted into individual empty wells of a 96-well plate using a BD FACSAria Fusion sorter. Plates were immediately sealed and stored at-80°C.

cDNA was generated from cells sorted using Superscript IV Reverse Transcriptase (Thermo Fisher), dNTPs (Thermo Fisher), random hexamers (Gene Link) and Ig gene-specific primers in a lysis buffer containing Igepal (Sigma), DTT and RNAseOUT (Thermo Fisher). Nested PCR amplification of heavy- and light-chain variable regions was performed using Hot Start DNA Polymerases (Qiagen, Thermo Fisher), and previously described primer sets [21,22]. Second round PCR primers were modified to include additional nucleotides overlapping with the expression vectors. PCR efficiency was assessed using 96w E-gels (Thermo Fisher). Paired wells picked individually, re-arrayed into new 96w plates and cloned in-frame into expression vectors encoding the human IgG1, Ig kappa or Ig lambda constant domains using the Gibson Assembly Enzyme mix (New England BioLabs) according to manufacturer instructions. Ligation reactions were transformed into DH5a competent E-coli, transferred into 1mL Plasmid+ media (Thomson Instrument Company) supplemented with antibiotic and grown overnight at 37C under agitation. The next day the cultures were used to inoculate duplicate cultures before being lysed for plasmid DNA extraction using NucleoSpin 96 miniprep kit (Macherey-Nagel, Takara). Cloned heavy- and light-chain variable regions were sequenced (Genewiz) and subsequently analyzed using the IMGT (International ImMunoGeneTics Information System, www.imgt.org) V-quest webserver [23].

<u>Flow cytometry-based cell surface ACE2 binding inhibition assay.</u>
Monoclonal antibody inhibition of SARS-CoV-2 S or RBD binding to cell surface hACE2 was performed by flow cytometry as follows. Purified mAbs were mixed with biotinylated SARS-CoV-2 S or RBD in the molar ratio of 4:1 on ice for 1h. In the meantime, HeLa-ACE2 cells were washed once with DPBS then detached by incubation with DPBS supplemented with 5 mM EDTA. The detached HeLa-ACE2 cells were washed and resuspended with FACS buffer (2% FBS and 1 mM EDTA in DPBS). 0.5 million Hela-ACE2 cells were added to mAb/S or RBD mixture and incubated at 4°C for 0.5 h. HeLa-ACE2 cells were then washed once with FACS buffer, resuspended FACS buffer with 1 µg/mL streptavidin-AF647 (Thermo, S21374) and incubated for another 0.5 h. After washing, HeLa-ACE2 cells were resuspended in FACS buffer in the presence of 2 µg/mL propidium iodide (Sigma, P4170-100MG) for live/dead staining. HeLa and HeLa-ACE2 cells stained with SARS-CoV-2 S or RBD alone were used as background and positive control separately. The AF647 mean fluorescence intensity (MFI) was determined from the gate of singlet and PI negative cells. The percentage of ACE2 binding inhibition was calculated using the following equation.

<u>Antibody expression and purification</u>
Antibodies HC and LC constructs were transiently expressed with the Expi293 Expression System (Thermo fisher). After 4 days, 24-deep well culture supernatants were harvested to be directly tested for binding and neutralization. Selected mAbs showing neutralizing activity in the HTP screening were re-expressed in small to medium scale cultures using individual colonies plasmid DNA, and IgG purified on Protein A sepharose (GE Healthcare).

<u>Epitope binning by bio-layer interferometry</u>
Neutralizing and non-neutralizing mAbs were binned into epitope specificities using an Octet RED384 system. 50-100 nM of HIS-tagged S or RBD protein antigens were captured using anti-Penta-HIS biosensors (18-5120, Molecular Devices). After antigen loading for 5 min, a saturating concentration of mAb (100 µg/mL) was added for 10 min. Competing concentrations of mAbs (25 µg/mL) were then added for 5 min to measure binding in the presence of saturating antibodies. All incubation steps were performed in 1x PBS.

<u>Surface Plasmon Resonance Methods</u>
SPR measurements were collected using a Biacore 8K instrument at 25°C. All experiments were carried out with a flow rate of 30 µL/min in a mobile phase of HBS-EP+ [0.01 M HEPES (pH 7.4), 0.15 M NaCl, 3 mM EDTA, 0.0005% (v/v) Surfactant P20]. Anti-Human IgG (Fc) antibody (Cytiva) was immobilized to

a density of ~7000-10000 RU via standard NHS/EDC coupling to a Series S CM-5 (Cytiva) sensor chip. A reference surface was generated through the same method.

For conventional kinetic/dose-response, listed antibodies were captured to ~50-100 RU via Fc-capture on the active flow cell prior to analyte injection. A concentration series of SARS-CoV-2 RBD was injected across the antibody and control surface for 2 min, followed by a 5 min dissociation phase using a multi-cycle method. Regeneration of the surface in between injections of SARS-CoV-2 RBD was achieved with a single, 120 s injection of 3M MgCl2. Kinetic analysis of each reference subtracted injection series was performed using the BIAEvaluation software (Cytiva). All sensorgram series were fit to a 1:1 (Langmuir) binding model of interaction.

A SPR assay was also used to assess the competition between SARS-CoV-2 RBD and ACE2 for binding to CC12.1 [24,25]. CC12.1 was captured to the surface of 3 flow cells to ~100 RU via Fc-capture. SARS-CoV-2 RBD was injected to each flow cell at a concentration of 50 nM to establish a basal level of SARS-CoV-2 RBD binding. This concentration was held constant for the competition experiments, which were carried out by varying the ACE2 concentration over eight points from 800 to 6.25 nM. To calculate residual SARS-CoV-2 RBD binding, the sensorgram responding to the corresponding ACE2 injection alone was subtracted from the SARS-CoV-2 RBD plus ACE2 injection series. The average response for the 5 s preceding the injection stop was plotted against the concentration of ACE2 and fit to a dose-response inhibition curve by nonlinear regression [log(inhibitor) vs. response – variable slope (4 parameters)] using GraphPad Prism. Regeneration between injections was carried out as noted above.

Animal Study

SARS-CoV-2 infection of 8-week old Syrian hamsters was achieved through the intranasal instillation of $10^6$ total PFU per animal in 100 µl of PBS. Animals weights were obtained during the study as a measure of disease progression. Treatment groups included the intraperitoneal injection of varying doses of monoclonal antibody. After 12-h, serum was obtained to quantify mAb titers and animals were infected as described above. At day-5 post-infection, lungs were harvested for analysis. Research protocol was approved and performed in accordance with Scripps Research IACUC Protocol #20-0003.

Viral load measurements

Viral RNA was isolated from lung tissue and subsequently amplified and quantified in a RT-qPCR reaction. Lung tissue was extracted at day 5 post infection. The lung tissue was divided into sections approximately 100-300mg in size. Samples were placed in 1mL of TRIzol-LS reagent (Invitrogen). Samples for virus load were then subjected to tissue homogenization using disposable pestles in 15mL conical tubes (Corning). Tissue homogenates were then spun down to remove any remaining cellular debris and the supernatant was added to a RNA purification column (Qiagen). Purified RNA was eluted in 80uL of DNase-, RNase-, endotoxin-free molecular biology grade water (Millipore) and quantified using a nanodrop (Thermo Fisher). RNA was then subjected to reverse transcription and quantitative PCR using the CDC's N1 primer sets (Forward 5'-GAC CCC AAA ATC AGC GAA AT-3'; Reverse 5'-TCT GGT TAC TGC CAG TTG AAT CTG-3') and a double-quenched (ZEN/Iowa Black FQ) and fluorescently labeled (FAM) probe (5'-FAM-ACC CCG CAT TAC GTT TGG TGG ACC-BHQ1-3') (Integrated DNA Technologies) on an BioRad CFX96 Real-Time instrument. For quantification, a standard curve was generated by diluting 1×1010 RNA copies SARS-CoV-2 genome/mL, 10-fold in water (Virapur). Every run utilized eight, tenfold serial dilutions of the standard. SARS-CoV-2-positive and -negative samples were included.

**Table S1. Cohort Demographics.**

| | PCR confirmed? | First blood collect date | Symptom onset (date) | Days post symptom onset | Age | Gender | Disease severity | Past medical history |
|---|---|---|---|---|---|---|---|---|
| **CC01** | Yes | 13-Feb | 9-Feb | 4 | 51 | F | Moderate | None |
| **CC04** | Yes | 16-Feb | 6-Feb | 10 | 56 | F | Mild | None |
| **CC05** | Yes | 14-Feb | 3-Feb | 11 | 53 | M | Moderate | None |
| **CC06** | Yes | 13-Mar | 5-Mar | 8 | 59 | M | Severe | Other |
| **CC08** | Yes | 23-Mar | 9-Mar | 14 | 62 | M | Severe | None |
| **CC09** | Yes | 26-Mar | 1-Mar | 25 | 44 | F | Mild | None |
| **CC10** | Yes | 26-Mar | 4-Mar | 22 | 45 | M | Mild-to-moderate | Asthma |
| **CC11** | Yes | 26-Mar | 1-Mar | 25 | 43 | F | Mild | None |
| **CC12** | Yes | 26-Mar | 1-Mar | 25 | 39 | F | Mild | None |
| **CC13** | Yes | 26-Mar | 1-Mar | 25 | 48 | F | Mild | None |
| **CC18** | Yes | 31-Mar | 8-Mar | 23 | 48 | F | Mild | None |
| **CC20** | No | 31-Mar | Unknown | - | 59 | M | Mild | None |
| **CC21** | Yes | 26-Mar | 15-Mar | 11 | 50 | M | Severe | HIV, HTN |
| **CC22** | Yes | 2-Apr | 27-Mar | 6 | 43 | M | Critical | None |
| **CC23** | Yes | 2-Apr | 15-Mar | 18 | 72 | F | Moderate | CKD, CAD |
| **CC24** | Yes | 7-Apr | 14-Mar | 24 | 32 | M | Mild | None |
| **CC25** | Yes | 7-Apr | 11-Mar | 27 | 42 | M | Mild | None |

Abbreviations: HLD, hyperlipidemia; HTN, hypertension; HCQ, hydroxychloroquine

**Table S2. Antibody sequence analysis.** Clonal families are highlighted in shaded colors.

| | Sequence characterisitics | | | |
|---|---|---|---|---|
| **mAb ID** | **VH Gene** | **% SHM (nt)** | **VL Gene** | **% SHM (nt)** |
| **CC12.1** | IGHV3-53*01 | 1.05 | IGKV1-9*01 | 1.08 |
| **CC12.2** | IGHV3-53*01 | 0.70 | IGKV3-20*01 | 3.55 |
| **CC12.3** | IGHV3-53*01 | 1.40 | IGKV3-20*01 | 0.36 |
| **CC12.4** | IGHV1-2*04 | 1.74 | IGLV2-8*01 | 1.74 |
| **CC12.5** | IGHV1-2*02 | 4.86 | IGLV2-14*01 | 3.47 |
| **CC12.6** | IGHV1-2*02 | 4.17 | IGLV2-14*01 | 1.74 |
| **CC12.7** | IGHV1-2*02 | 7.99 | IGLV2-14*01 | 6.60 |
| **CC12.8** | IGHV1-2*02 | 3.12 | IGLV2-14*01 | 1.04 |
| **CC12.9** | IGHV1-2*02 | 3.47 | IGLV2-14*01 | 1.39 |
| **CC12.10** | IGHV1-2*02 | 4.51 | IGLV2-14*01 | 4.17 |
| **CC12.11** | IGHV1-2*02 | 3.47 | IGLV2-14*01 | 1.74 |
| **CC12.12** | IGHV1-2*02 | 3.47 | IGLV2-14*01 | 2.08 |
| **CC12.13** | IGHV3-53*01 | 0.35 | IGKV1-33*01 | 3.23 |
| **CC12.14** | IGHV3-21*01 | 3.47 | IGKV2-30*01 | 1.70 |
| **CC12.15** | IGHV3-48*03 | 1.04 | IGLV1-40*01 | 1.04 |
| **CC12.16** | IGHV3-33*01 | 1.74 | IGLV3-21*02 | 1.08 |
| **CC12.17** | IGHV3-30*03 | 1.39 | IGLV3-21*03 | 1.79 |
| **CC12.18** | IGHV1-46*01 | 0.00 | IGLV6-57*01 | 1.03 |
| **CC12.19** | IGHV3-23*04 | 1.04 | IGLV3-21*02 | 1.79 |
| **CC12.20** | IGHV3-30*03 | 1.04 | IGLV1-47*01 | 1.05 |
| **CC12.21** | IGHV1-24*01 | 1.39 | IGLV1-44*01 | 0.70 |
| **CC12.22** | IGHV1-24*01 | 1.04 | IGKV3-15*01 | 0.00 |
| **CC12.23** | IGHV4-39*01 | 0.69 | IGLV3-25*03 | 0.36 |
| **CC12.24** | IGHV3-30*03 | 1.04 | IGKV1-39*01 | 0.72 |
| **CC12.25** | IGHV3-23*04 | 0.35 | IGLV1-44*01 | 2.46 |
| **CC12.26** | IGHV5-51*01 | 0.35 | IGLV7-43*01 | 1.04 |
| **CC12.27** | IGHV1-2*02 | 1.04 | IGLV2-23*01 | 0.35 |

### Table S3. Hamster protection study summary.

| Animal ID | CC12.1 Dose (μg) | Weight (g) at Day 0 | Weight (g) at Day 1 | Weight (g) at Day 3 | Weight (g) at Day 4 | Weight (g) at Day 5 | Weight (g) at Day 6 | Lung sample mass (mg) | Virus load (absolute) | Virus load per mg (VL/mg) | CC2.1 serum titer at Day 0 (μg/mL) | CC2.1 serum titer at Day 5 (μg/mL) |
|---|---|---|---|---|---|---|---|---|---|---|---|---|
| **1-334** | Ctrl | 124 | 120 | 114 | 108 | 106 | | 132 | 47381442133 | 358950319 | 0.00 | 0 |
| **1-335** | Ctrl | 109 | 100 | 98 | 94 | 93 | | 166.6 | 2079731197 | 12483381 | 0.00 | 0 |
| **1-338** | Ctrl | 122 | 121 | 116 | 111 | 110 | | 98.1 | 3929842857 | 40059560 | 0.00 | 0 |
| **1-432** | Ctrl | 131 | 126 | 120 | 114 | 112 | | 248.4 | 952477443 | 3834450 | 0.00 | 0 |
| **1-436** | Ctrl | 120 | 118 | 108 | 103 | 102 | | 202.8 | 1097002828 | 5409284 | 0.00 | 0 |
| **1-433** | Ctrl | 120 | 118 | 110 | 108 | 104 | | 144.2 | 29037705468 | 201371050 | 0.00 | 0 |
| **1-440** | 2000 | 130 | 122 | 145 | 142 | 141 | | 139.2 | 22203736 | 159510 | 46.99 | 97 |
| **1-451** | 2000 | 134 | 128 | 130 | 135 | 138 | | 137.8 | 23367181 | 169573 | 28.46 | 68 |
| **1-437** | 2000 | 124 | 117 | 128 | 132 | 131 | | 77.7 | 1758266 | 22629 | 20.77 | 135 |
| **1-447** | 2000 | 109 | 100 | 110 | 114 | 118 | | 180 | 59773323 | 332074 | 49.67 | 0 |
| **1-449** | 2000 | 108 | 100 | 102 | 104 | 104 | | 163.8 | 7562491 | 46169 | 29.95 | 75 |
| **1-448** | 2000 | 120 | 110 | 118 | 122 | 124 | | 134.7 | 8504808 | 63139 | 29.36 | 98 |
| **1-443** | 500 | 119 | 110 | 120 | 121 | 123 | | 39.3 | 7908498 | 201234 | 27.19 | 31 |
| **1-427** | 500 | 143 | 131 | 144 | 146 | 150 | | 58.2 | 1731959 | 29759 | 19.92 | 42 |
| **1-431** | 500 | 120 | 110 | 112 | 114 | 116 | | 155.7 | 111573372 | 716592 | 12.38 | 63 |
| **1-429** | 500 | 119 | 110 | 118 | 120 | 120 | | 56.5 | 2427923 | 42972 | 18.89 | 42 |
| **1-428** | 500 | 98 | 90 | 94 | 96 | 98 | | 66.3 | 12872027 | 194148 | 26.27 | 49 |
| **1-441** | 500 | 122 | 116 | 120 | 121 | 124 | | 164.4 | 17194913 | 104592 | 38.80 | 42 |
| **1-452** | 125 | 130 | 130 | 126 | 122 | 121 | | 135.7 | 9227879 | 68002 | 28.42 | 1 |
| **1-445** | 125 | 120 | 110 | 106 | 106 | 104 | | 145.5 | 74616429 | 512828 | 15.04 | 1 |
| **1-456** | 125 | 90 | 92 | 90 | 88 | 87 | | 135.4 | 96077317 | 709581 | 14.29 | 2 |
| **1-454** | 125 | 130 | 122 | 124 | 123 | 124 | | 142.6 | 8874684 | 62235 | 5.16 | 2 |
| **1-453** | 125 | 128 | 120 | 115 | 114 | 118 | | 188.4 | 2369242 | 12576 | 6.83 | 3 |
| **1-455** | 125 | 120 | 111 | 108 | 104 | 104 | | 165.9 | 11500369 | 69321 | 4.20 | 6 |
| **1-442** | 31 | 110 | 100 | 98 | 95 | 94 | | 113.3 | 146999226 | 1297434 | 9.82 | 0 |
| **1-454** | 31 | 118 | 102 | 106 | 106 | 103 | | 60.8 | 230749502 | 3795222 | 3.50 | 0 |
| **1-446** | 31 | 110 | 100 | 98 | 96 | 94 | | 145.8 | 7505018 | 51475 | 9.49 | 0 |
| **1-443** | 31 | 119 | 110 | 107 | 102 | 101 | | 50 | 1264877867 | 25297557 | 6.67 | 0 |
| **1-444** | 31 | 120 | 101 | 100 | 97 | 94 | | 65.8 | 385355044 | 5856460 | 1.44 | 0 |
| **1-445** | 31 | 100 | 90 | 86 | 84 | 84 | | 65.2 | 1077752020 | 16529939 | 25.30 | 0 |
| **1-458** | 8 | 127 | 110 | 112 | 103 | 102 | | 126.6 | 247726739 | 1956767 | 0.11 | 0 |
| **1-459** | 8 | 120 | 111 | 100 | 94 | 96 | | 219.1 | 368023811 | 1679707 | 0.00 | 0 |
| **1-457** | 8 | 118 | 110 | 103 | 100 | 97 | | 244.3 | 16935521 | 69323 | 0.29 | 0 |
| **1-460** | 8 | 100 | 96 | 92 | 87 | 85 | | 104.4 | 1153435020 | 11048228 | 0.06 | 0 |
| **1-463** | 8 | 110 | 102 | 97 | 92 | 94 | | 143.9 | 822113391 | 5713088 | 0.00 | 0 |
| **1-461** | 8 | 120 | 115 | 112 | 108 | 104 | | 83 | 2813485933 | 33897421 | 0.12 | 0 |

| Animal | CC12.26 Dose (μg) | Weight (g) at Day 0 | Weight (g) at Day 1 | Weight (g) at Day 3 | Weight (g) at Day 4 | Weight (g) at Day 5 | Weight (g) at Day 6 | Lung sample mass (mg) | Virus load (absolute) | Virus load per mg (VL/mg) | CC2.26 serum titer at Day 0 (μg/mL) |
|---|---|---|---|---|---|---|---|---|---|---|---|
| **26-422** | Ctrl | 103 | | 100 | 97 | | 96 | | | | 0.00 |
| **26-410** | Ctrl | 110 | | 100 | 96 | | 93 | | | | 0.00 |
| **26-425** | Ctrl | 120 | | 110 | 110 | | 102 | | | | 0.00 |
| **26-423** | Ctrl | 112 | | 110 | 104 | | 105 | | | | 0.00 |
| **26-424** | Ctrl | 90 | | 88 | 82 | | 78 | | | | 0.00 |
| **26-426** | Ctrl | 108 | | 100 | 100 | | 93 | | | | 0.00 |
| **26-414** | 2000 | 100 | | 100 | 100 | | 96 | | | | 94.69 |
| **26-415** | 2000 | 130 | | 126 | 126 | | 123 | | | | 88.06 |
| **26-408** | 2000 | 128 | | 126 | 121 | | 119 | | | | 88.13 |
| **26-416** | 2000 | 122 | | 118 | 110 | | 110 | | | | 105.13 |
| **26-413** | 2000 | 120 | | 118 | 118 | | 112 | | | | 70.32 |
| **26-412** | 2000 | 110 | | 100 | 98 | | 96 | | | | 34.93 |
| **26-409** | 500 | 120 | | 118 | 106 | | 102 | | | | 38.02 |
| **26-403** | 500 | 120 | | 118 | 110 | | 110 | | | | 34.76 |
| **26-404** | 500 | 130 | | 120 | 110 | | 104 | | | | 28.39 |
| **26-402** | 500 | 110 | | 102 | 100 | | 101 | | | | 42.84 |
| **26-406** | 500 | 110 | | 114 | 100 | | 100 | | | | 19.64 |
| **26-405** | 500 | 120 | | 110 | 109 | | 110 | | | | 0.34 |
| **26-388** | 125 | 100 | | 100 | 108 | | 110 | | | | 6.83 |
| **26-390** | 125 | 120 | | 119 | 112 | | 108 | | | | 6.01 |
| **26-391** | 125 | 119 | | 116 | 110 | | 107 | | | | 7.01 |
| **26-439** | 125 | 100 | | 100 | 96 | | 93 | | | | 6.58 |
| **26-389** | 125 | 110 | | 104 | 102 | | 108 | | | | 4.46 |
| **26-387** | 125 | 138 | | 130 | 123 | | 118 | | | | 0.00 |
| **26-411** | 31 | 124 | | 121 | 110 | | 108 | | | | 2.26 |
| **26-417** | 31 | 100 | | 98 | 92 | | 90 | | | | 2.30 |
| **26-421** | 31 | 100 | | 100 | 94 | | 95 | | | | 0.00 |
| **26-419** | 31 | 100 | | 100 | 92 | | 89 | | | | 2.08 |
| **26-420** | 31 | 112 | | 110 | 104 | | 103 | | | | 0.83 |
| **26-418** | 31 | 133 | | 132 | 120 | | 115 | | | | 1.17 |
| **26-407** | 8 | 120 | | 110 | 110 | | 107 | | | | 0.00 |
| **26-393** | 8 | 101 | | 100 | 98 | | 96 | | | | 0.25 |
| **26-394** | 8 | 100 | | 99 | 90 | | 87 | | | | 0.35 |
| **26-396** | 8 | 118 | | 110 | 108 | | 105 | | | | 0.00 |
| **26-395** | 8 | 123 | | 122 | 118 | | 114 | | | | 0.13 |
| **26-392** | 8 | 124 | | 120 | 110 | | 106 | | | | 0.16 |

**Table S4. Hamster protection study statistics.** Each group was compared to the Den3 IgG control group

| | | | CC12.1 protection experiment | | | | | | | | | | | | | |
|---|---|---|---|---|---|---|---|---|---|---|---|---|---|---|---|---|
| | | | IgG Day 0 | | Weigh Day 0 | | Weigh Day 1 | | Weigh Day 3 | | Weigh Day 4 | | Weigh Day 5 | | Viral Load Day 5 | |
| Statistical test | | Ab Dose (µg) | Summary | Adjusted P-Value | Summary | Adjusted P-Value | Summary | Adjusted P-Value | Summary | Adjusted P-Value | Summary | Adjusted P-Value | Summary | Adjusted P-Value | Summary | Adjusted P-Value |
| Grouped Parametric | One-way ANOVA | 2000 | **** | <0.0001 | ns | 0.9957 | ns | 0.0972 | ** | 0.0014 | **** | <0.0001 | **** | <0.0001 | * | 0.0274 |
| | | 500 | **** | <0.0001 | ns | 0.9957 | ns | 0.0704 | * | 0.0319 | **** | <0.0001 | **** | <0.0001 | * | 0.0274 |
| | | 125 | * | 0.0318 | ns | 0.9957 | ns | 0.5244 | ns | 0.4496 | ns | 0.2054 | * | 0.0269 | * | 0.0274 |
| | | 31 | ns | 0.0923 | ns | 0.7037 | *** | 0.0005 | ns | 0.2663 | ns | 0.3002 | ns | 0.2835 | * | 0.0274 |
| | | 8 | ns | 0.9834 | ns | 0.8946 | ns | 0.0795 | ns | 0.3494 | ns | 0.2054 | ns | 0.2309 | * | 0.0274 |
| Grouped Non-Parametric | Kruskall-Wallis | 2000 | **** | <0.0001 | ns | >0.9999 | ns | 0.6064 | ns | 0.1275 | * | 0.0192 | * | 0.0128 | ** | 0.0025 |
| | | 500 | ** | 0.0011 | ns | >0.9999 | ns | 0.0884 | ns | 0.2931 | ns | 0.0709 | * | 0.0477 | ** | 0.0074 |
| | | 125 | * | 0.0377 | ns | >0.9999 | ns | >0.9999 | ns | >0.9999 | ns | >0.9999 | ns | >0.9999 | ** | 0.0041 |
| | | 31 | ns | 0.1111 | ns | 0.33 | *** | 0.0007 | ns | 0.6242 | ns | >0.9999 | ns | >0.9999 | ns | >0.9999 |
| | | 8 | ns | >0.9999 | ns | >0.9999 | ns | 0.7306 | ns | >0.9999 | ns | >0.9999 | ns | >0.9999 | ns | >0.9999 |
| Individual Parametric | t-test | 2000 | **** | <0.0001 | ns | 0.9753 | * | 0.021 | ** | 0.0051 | **** | <0.0001 | **** | <0.0001 | ns | 0.1131 |
| | | 500 | **** | <0.0001 | ns | 0.9008 | ** | 0.006 | ** | 0.0012 | **** | <0.0001 | **** | <0.0001 | ns | 0.1134 |
| | | 125 | ** | 0.008 | ns | 0.8502 | ns | 0.6122 | ns | 0.4247 | ns | 0.0784 | * | 0.0179 | ns | 0.1135 |
| | | 31 | * | 0.0219 | ns | 0.0863 | ** | 0.0012 | * | 0.0163 | ns | 0.1784 | ns | 0.185 | ns | 0.1435 |
| | | 8 | ns | 0.0541 | ns | 0.3115 | ns | 0.0504 | ns | 0.1005 | ns | 0.0784 | ns | 0.0524 | ns | 0.145 |
| Individual Non-Parametric | Mann-Whitney | 2000 | ** | 0.0022 | ns | >0.9999 | * | 0.0303 | ** | 0.0043 | ** | 0.0022 | ** | 0.0022 | ** | 0.0022 |
| | | 500 | ** | 0.0022 | ns | 0.5152 | * | 0.0238 | ** | 0.0043 | ** | 0.0022 | ** | 0.0022 | ** | 0.0022 |
| | | 125 | ** | 0.0022 | ns | 0.9264 | ns | 0.6667 | ns | 0.8506 | ns | 0.132 | * | 0.0152 | ** | 0.0022 |
| | | 31 | ** | 0.0022 | ns | 0.0519 | ** | 0.0043 | ** | 0.0043 | ns | 0.2403 | ns | 0.1688 | ns | 0.132 |
| | | 8 | ns | 0.0606 | ns | 0.29 | * | 0.039 | ns | 0.2403 | ns | 0.0693 | ns | 0.0866 | ns | 0.0931 |

| | | | CC12.23 protection experiment | | | | | | | | | | |
|---|---|---|---|---|---|---|---|---|---|---|---|---|---|
| | | | IgG Day 0 | | Weigh Day 0 | | | Weigh Day 3 | | Weigh Day 4 | | Weigh Day 5 | |
| | | Ab Dose (µg) | Summary | Adjusted P-Value | Summary | Adjusted P-Value | | Summary | Adjusted P-Value | Summary | Adjusted P-Value | Summary | Adjusted P-Value |
| Grouped Parametric | One-way ANOVA | 2000 | **** | <0.0001 | ns | 0.4367 | | ns | 0.5798 | ns | 0.5035 | ns | 0.4976 |
| | | 500 | ** | 0.0018 | ns | 0.4367 | | ns | 0.5798 | ns | 0.7427 | ns | 0.9187 |
| | | 125 | ns | 0.8462 | ns | 0.6359 | | ns | 0.4409 | ns | 0.4521 | ns | 0.1817 |
| | | 31 | ns | 0.9735 | ns | 0.6359 | | ns | 0.1595 | ns | 0.9931 | ns | 0.9139 |
| | | 8 | ns | 0.9831 | ns | 0.6359 | | ns | 0.5798 | ns | 0.9115 | ns | 0.9139 |
| Grouped Non-Parametric | Kruskall-Wallis | 2000 | **** | <0.0001 | ns | 0.5146 | | ns | >0.9999 | ns | >0.9999 | ns | 0.2503 |
| | | 500 | ** | 0.0033 | ns | 0.6082 | | ns | >0.9999 | ns | 0.8312 | ns | >0.9999 |
| | | 125 | ns | 0.0769 | ns | >0.9999 | | ns | 0.3718 | ns | >0.9999 | ns | 0.5597 |
| | | 31 | ns | 0.4173 | ns | >0.9999 | | ns | 0.0878 | ns | >0.9999 | ns | >0.9999 |
| | | 8 | ns | >0.9999 | ns | >0.9999 | | ns | >0.9999 | ns | >0.9999 | ns | >0.9999 |
| Individual Parametric | t-test | 2000 | **** | <0.0001 | ns | 0.1028 | | ns | 0.2721 | ns | 0.1468 | ns | 0.0785 |
| | | 500 | ** | 0.0015 | ns | 0.0549 | | ns | 0.5508 | ns | 0.1866 | ns | 0.9123 |
| | | 125 | *** | 0.0008 | ns | 0.332 | | ns | 0.1307 | ns | 0.2308 | ns | 0.1449 |
| | | 31 | ** | 0.0036 | ns | 0.5568 | | * | 0.0159 | ns | 0.9877 | ns | 0.4424 |
| | | 8 | ns | 0.2431 | ns | 0.2652 | | ns | 0.3764 | ns | 0.6137 | ns | 0.5128 |
| Individual Non-Parametric | Mann-Whitney | 2000 | ** | 0.0022 | ns | 0.119 | | ns | 0.329 | ns | 0.2403 | ns | 0.0931 |
| | | 500 | ** | 0.0022 | ns | 0.0758 | | ns | 0.3268 | ns | 0.0693 | ns | 0.9848 |
| | | 125 | * | 0.0152 | ns | 0.5974 | | ns | 0.1255 | ns | 0.3095 | ns | 0.2403 |
| | | 31 | * | 0.0152 | ns | 0.8463 | | * | 0.0173 | ns | 0.9675 | ns | 0.3095 |
| | | 8 | ns | 0.0606 | ns | 0.329 | | ns | 0.2576 | ns | 0.9654 | ns | 0.3939 |

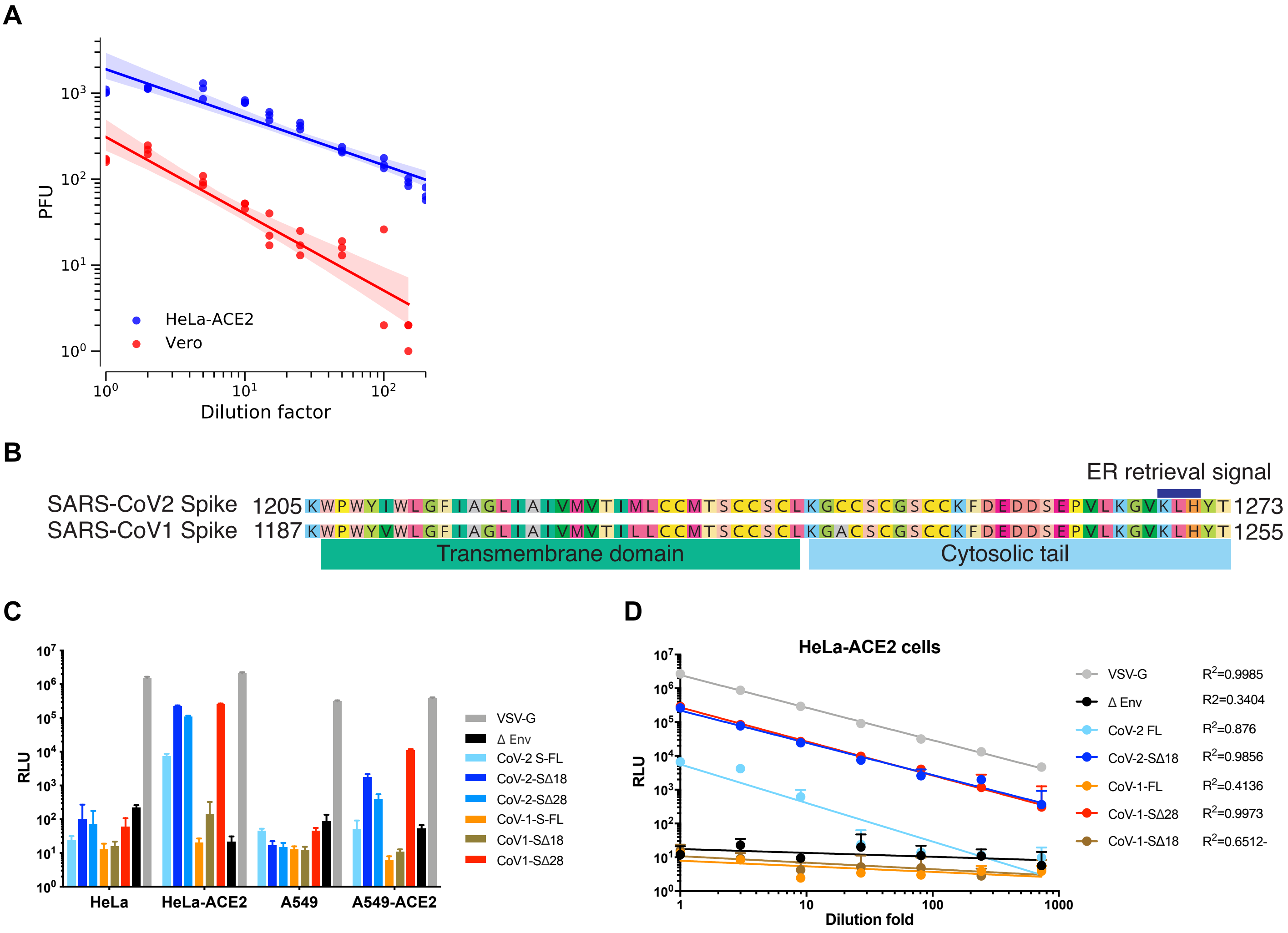

**Fig. S1. Neutralization assay development.** (**A**) Vero and HeLa-ACE2 cells were infected with serially diluted SARS-CoV-2. Both cell types were plated at 1000 cells/well. The HeLa-ACE2 cell line showed 100% infection at a dilution factor of 2. (**B**) The alignment of SARS-CoV-1 and SARS-CoV-2 transmembrane domain and cytoplasmic tail. (**C**) Comparison of the infectious efficiency of MLV viral particles pseudotyped with the indicated spike (S) proteins: FL indicates full-length spike, D18 and D28 denote C-terminal truncations. Also included were control VSV-G-pseudotyped virions and empty vector control (D Env). Target cells were HeLa cells, A549 cells, HeLa or A549 cells overexpressing human ACE2. After 48h of infection, luciferase expression was assessed. (**D**) Pseudovirions were titrated in HeLa-ACE2 cells in 3-fold serial dilutions. Relative luciferase and fold-dilution are plotted as indicated. Correlation coefficient ($R^2$) was calculated using linear regression in Prism.

## A

| Participant ID | ELISA Binding $ED_{50}$ | | | | | Cell Surface Binding $ID_{50}$ (%Positive Cells) | | Pseudovirus Neutralization $ID_{50}$ | | | Live Virus Neut. $ID_{50}$ |
|---|---|---|---|---|---|---|---|---|---|---|---|
| | SARS-CoV-2 S | SARS-CoV-2 RBD | BSA | SARS-CoV-1 S | SARS-CoV-1 RBD | SARS-CoV-2 Spike | SARS-CoV-1 Spike | SARS-CoV-2 | SARS-CoV-1 | VSV-G | SARS-CoV-2 |
| CC01 | 22 | <20 | <20 | <20 | <20 | <30 | 75 | 49 | <20 | <20 | <20 |
| CC04 | 17036 | 2233 | <20 | 791 | 100 | 29971 | 4945 | 2320 | 90 | <20 | 3382 |
| CC05 | <20 | <20 | <20 | <20 | <20 | <30 | <30 | <20 | <20 | <20 | <20 |
| CC06 | 11188 | 1406 | <20 | 1095 | 39 | 297 | 285 | 1509 | <20 | <20 | 122 |
| CC08 | 221 | 20 | <20 | <20 | <20 | 152 | 368 | <20 | <20 | <20 | 35 |
| CC09 | 83 | 28 | <20 | 22 | <20 | 158 | 251 | 38 | <20 | <20 | 55 |
| CC10 | 552 | 110 | <20 | 57 | <20 | 281 | 334 | 196 | <20 | <20 | 538 |
| CC11 | 113 | 26 | <20 | 34 | <20 | 395 | 464 | 85 | <20 | <20 | 108 |
| CC12 | 307 | 24 | <20 | 28 | <20 | 572 | 326 | 630 | <20 | <20 | 478 |
| CC13 | 117 | 29 | <20 | 20 | <20 | 378 | 597 | 38 | <20 | <20 | 122 |
| CC18 | 41 | 23 | <20 | <20 | <20 | 52 | 102 | 23 | <20 | <20 | 43 |
| CC20 | <20 | <20 | <20 | <20 | <20 | <30 | 101 | <20 | <20 | <20 | <20 |
| CC21 | 3064 | 614 | <20 | 434 | 192 | 3440 | 1697 | 1941 | <20 | <20 | 1581 |
| CC22 | 1202 | 135 | <20 | 101 | <20 | 2166 | 1161 | 406 | <20 | <20 | 1478 |
| CC23 | 186 | <20 | <20 | 89 | <20 | 282 | 416 | 32 | <20 | <20 | 105 |
| CC24 | 343 | 51 | <20 | 32 | 22 | 458 | 211 | 24 | <20 | <20 | 106 |
| CC25 | 1386 | 560 | <20 | 300 | 42 | 1490 | 668 | 702 | <20 | <20 | 2522 |

## B

| | SARS-CoV-2 (Pseudotyped) | | SARS-CoV-2 (Live Virus) | |
|---|---|---|---|---|
| **ID50** | # of Donors | % of Donors | # of Donors | % of Donors |
| **<500** | 12 | 70 | 12 | 70 |
| **500-999** | 2 | 12 | 1 | 6 |
| **1000-2500** | 3 | 18 | 2 | 12 |
| **>2500** | 0 | 0 | 2 | 12 |

## C

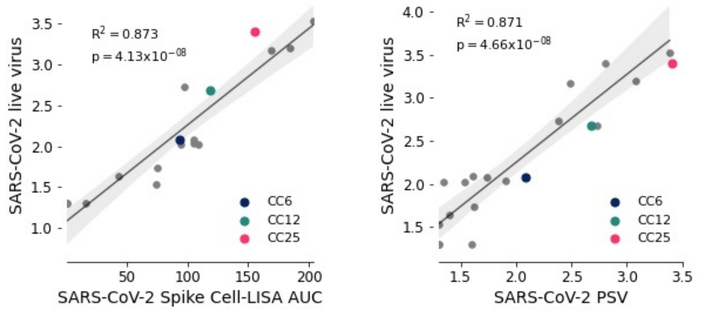

**Fig. S2. Evaluation of SARS-2 neutralizing antibody responses in CC participants.** Plasma samples collected from 17 COVID-19 Cohort participants were tested for SARS-CoV-1 and CoV-2 pseudotyped and live replicating virus neutralization, cell surface binding (Cell-LISA) to Spike protein and ELISA binding to recombinant S and RBD proteins. (**A**) Plasma titers are indicated. VSV-G was used as negative control. (**B**) Prevalence of potent and weak serum nAb responses to SARS-CoV-2 is compared for pseudotyped and live virus assays for 17 participants, 15 days post symptom onset. (**C)** Correlation between live virus neutralization and Cell-LISA binding assay (left) or pseudovirus neutralization (right) of participant plasma. For both neutralization assays, $IC_{50}$ values are $log_{10}$ transformed before plotting. Linear regression lines are shown. The 95% confidence interval of the regression line is shown in light grey shading and was computed using 1,000 bootstrap resamplings. $R^2$ and p values are indicated. AUC: Area-under-the-curve. Donors from whom mAbs were isolated are specifically highlighted in dark blue (CC6), pine green (CC12) and hot pink (CC25).

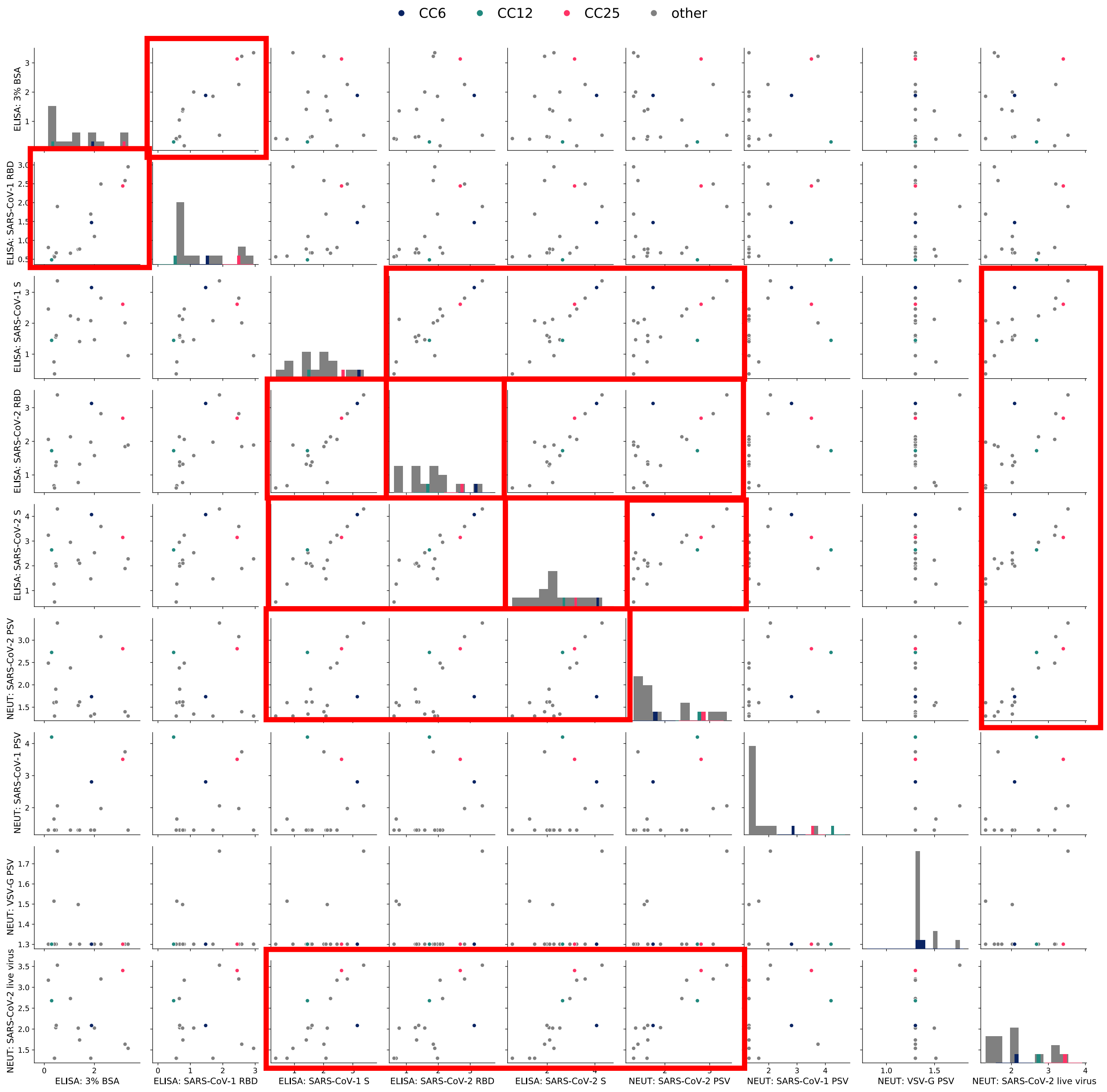

**Fig. S3. Evaluation of SARS-2 neutralizing antibody responses in CC participants.** Full correlation matrix of individual parameters used to evaluate plasma Ab responses.

**A**

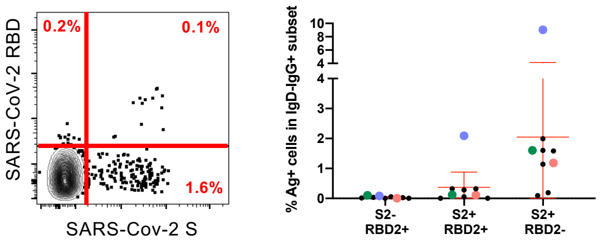

**B**

| Sort | Plate | # wells | Sample#1 | Sample#2 | HC Global | KC Global | LC Global | PAIRS | HC Cloned | KC Cloned | LC Cloned | #HKL | mAbs Transfected |
|---|---|---|---|---|---|---|---|---|---|---|---|---|---|
| | 1 | 96 | CC7 | CC9 | 59% | 36% | 40% | 44% | | | | | |
| | 2 | 96 | | CC9 | 77% | 56% | 38% | 64% | | | | | |
| | 3 | 96 | | CC9 | 76% | 46% | 36% | 52% | 287 | 144 | 101 | 38 | 287 |
| | 4 | 96 | | CC9 | 80% | 48% | 40% | 59% | | | | | |
| | 5 | 96 | | CC9 | 77% | 58% | 24% | 64% | | | | | |
| | 6 | 96 | CC10 | CC9 | 93% | 53% | 39% | 80% | 39 | 15 | 15 | 2 | 29 |
| | 10 | 96 | CC12 | CC11 | 69% | 39% | 50% | 66% | | | | | |
| | 11 | 96 | CC12 | | 83% | 50% | 41% | 73% | | | | | |
| Sort#1 | 12 | 96 | CC12 | | 75% | 43% | 46% | 74% | 273 | 154 | 107 | 28 | 273 |
| | 13 | 96 | CC12 | | 74% | 45% | 46% | 72% | | | | | |
| | 14 | 96 | CC12 | CC13 | 73% | 42% | 44% | 71% | | | | | |
| | 15 | 96 | | CC13 | 85% | 36% | 51% | 74% | | | | | |
| | 7 | 96 | CC10 | | 71% | 34% | 38% | 47% | | | | | |
| | 8 | 96 | | CC11 | 79% | 42% | 46% | 77% | | | | | |
| | 9 | 96 | | CC11 | 72% | 39% | 44% | 67% | | | | | |
| | 16 | 96 | | CC13 | 80% | 44% | 51% | 72% | 586 | 322 | 223 | 55 | 586 |
| | 17 | 88 | | CC13 | 73% | 38% | 32% | 63% | | | | | |
| | 18 | 96 | CoVID19- | | 55% | 44% | 20% | 39% | | | | | |
| | 1 | 96 | | CC12 | 70% | 32% | 41% | 58% | | | | | |
| Sort#2 | 2 | 96 | | CC12 | 81% | 43% | 48% | 74% | 183 | 96 | 106 | 16 | 183 |
| | 3 | 96 | | CC12 | 69% | 39% | 33% | 58% | | | | | |
| | 1 | 96 | CC6 | | 61% | 28% | 41% | 57% | | | | | |
| | 2 | 96 | CC6 | | 63% | 31% | 46% | 57% | | | | | |
| Sort#3 | 3 | 96 | CC6 | | 67% | 45% | 32% | 58% | 242 | 117 | 145 | 20 | 242 |
| | 4 | 96 | CC6 | | 69% | 26% | 47% | 60% | | | | | |
| | 5 | 96 | CC6 | | 61% | 42% | 36% | 55% | | | | | |
| | 1 | 96 | CC25 | | 72% | 48% | 43% | 66% | | | | | |
| | 2 | 96 | CC25 | | 77% | 50% | 41% | 70% | | | | | |
| | 3 | 96 | CC25 | | 77% | 49% | 38% | 65% | | | | | |
| Sort#4 | 4 | 96 | CC25 | CC6 | 80% | 52% | 40% | 75% | 445 | 273 | 96 | 58 | 445 |
| | 5 | 96 | | CC6 | 79% | 47% | 49% | 69% | | | | | |
| | 6 | 96 | | CC6 | 78% | 48% | 41% | 70% | | | | | |
| | 7 | 96 | | CC6 | 82% | 44% | 56% | 77% | | | | | |
| **TOTAL** | 39 | 3160 | 8 | | 2251 | 1301 | 1248 | 2055 | 2055 | 1121 | 793 | 217 | 2045 |

**C**

| | Heavy Chain (H) Colonies | Kappa chain (K) Colonies | Lambda chain (L) Colonies | H+K+L All Colonies |
|---|---|---|---|---|
| Total Colonies Screened | 819 | 819 | 819 | 2457 |
| Off-frame insert | 2 | 2 | 0 | 0.2% |
| IMGT: No Results | 45 | 4 | 9 | 2.4% |
| IMGT: Unproductive | 33 | 20 | 12 | 2.6% |
| IMGT: No rearrangement found | 7 | 0 | 0 | 0.3% |
| Total Cloning Reactions | 273 | 273 | 273 | 819 |
| Overall Cloning Failures | 12 | 18 | 11 | 5.0% |
| Total Cloned Pairs | | 273 | | 273 |
| Predicted Unproductive Pairs | | 37 | | 13.6% |

**Fig. S4. SARS-CoV-2 mAb isolation, cloning and expression summary. (A)** FACS frequency of memory (CD3-CD19+IgD-IgG+) B cells binding to SARS-CoV-2 S-protein (S2+) and or RBD subunit (RBD2+) in COVID-19 individuals. (Left) A representative fluorescence dot plot is shown for CC12. (Right) Frequencies for all donors are plotted. Average and SD are indicated with red lines. (**B**) The following statistics are tabulated **:** number of SARS-CoV-2 specific B-cells sorted for each donor in each experiment, efficiency of heavy and light chains PCR recovery (% of PCR positive wells, color gradient with warmer color indicating higher efficiency), number of heavy and light chains amplicons cloned; number of and H+L pairs transfected (green) for high-throughput functional screening. (**C**) Three individual colonies were picked from 273 paired H+L cloning reactions, send for sequencing and analyzed using IMGT V-Quest. Cloning efficiency statistics are indicated.

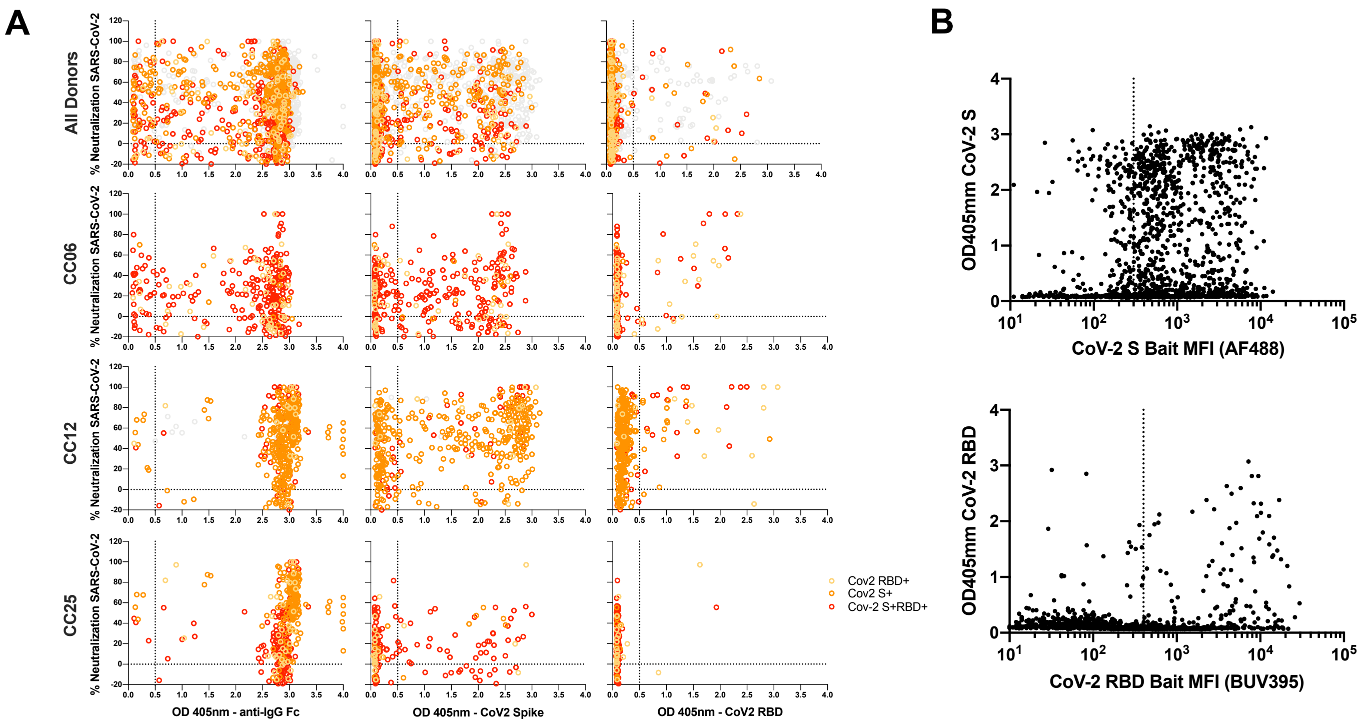

**Fig. S5. Functional screening of Ab H+L pairs rescued from SARS-CoV-2-specific single B-cell sorting.** (**A**) Cloned H+L chain pairs isolated from SARS-2 specific single B-cells were transfected into a high efficiency expression cell line. Small-scale culture supernatants were harvested at day5-post transfection and evaluated for the presence of IgG, binding to recombinant SARS-CoV-2 S-protein and RBD subunit as well as for pseudotyped SARS-CoV-2 neutralization. ELISA considered positive when OD405nm was > 0.5 (dotted line). Results are plotted to show the proportion of expressed, binding and neutralizing pairs. (**B**) Correlation between ELISA binding signal (OD 405nm) and corresponding sorted cell staining level (MFI) for each antigenic bait (SARS-CoV-2 S-protein or RBD).

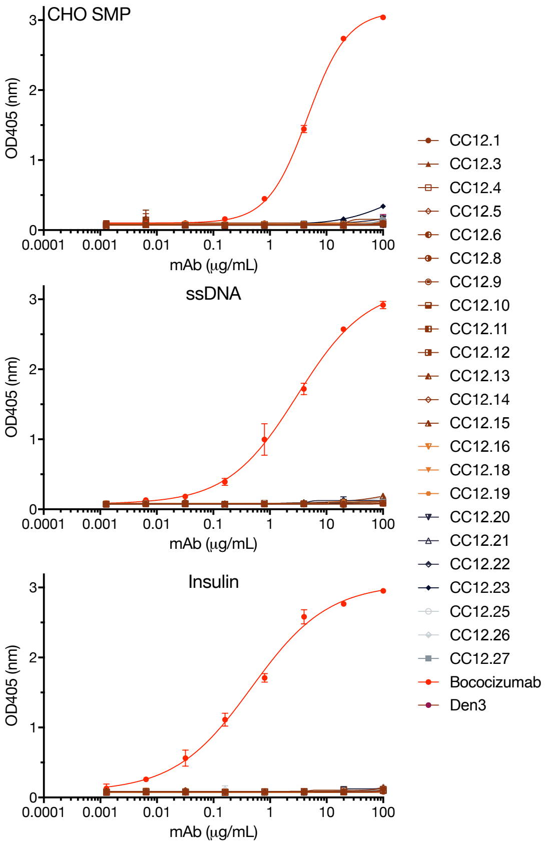

**Fig S6. Evaluation of SARS-CoV-2 specific mAbs for polyreactivity and autoreactivity.** Antibodies were tested by ELISA for binding to several polyspecificity reagents (PSR): CHO-cell soluble membrane protein extracts (SMP), single stranded DNA (ssDNA) and Insulin.

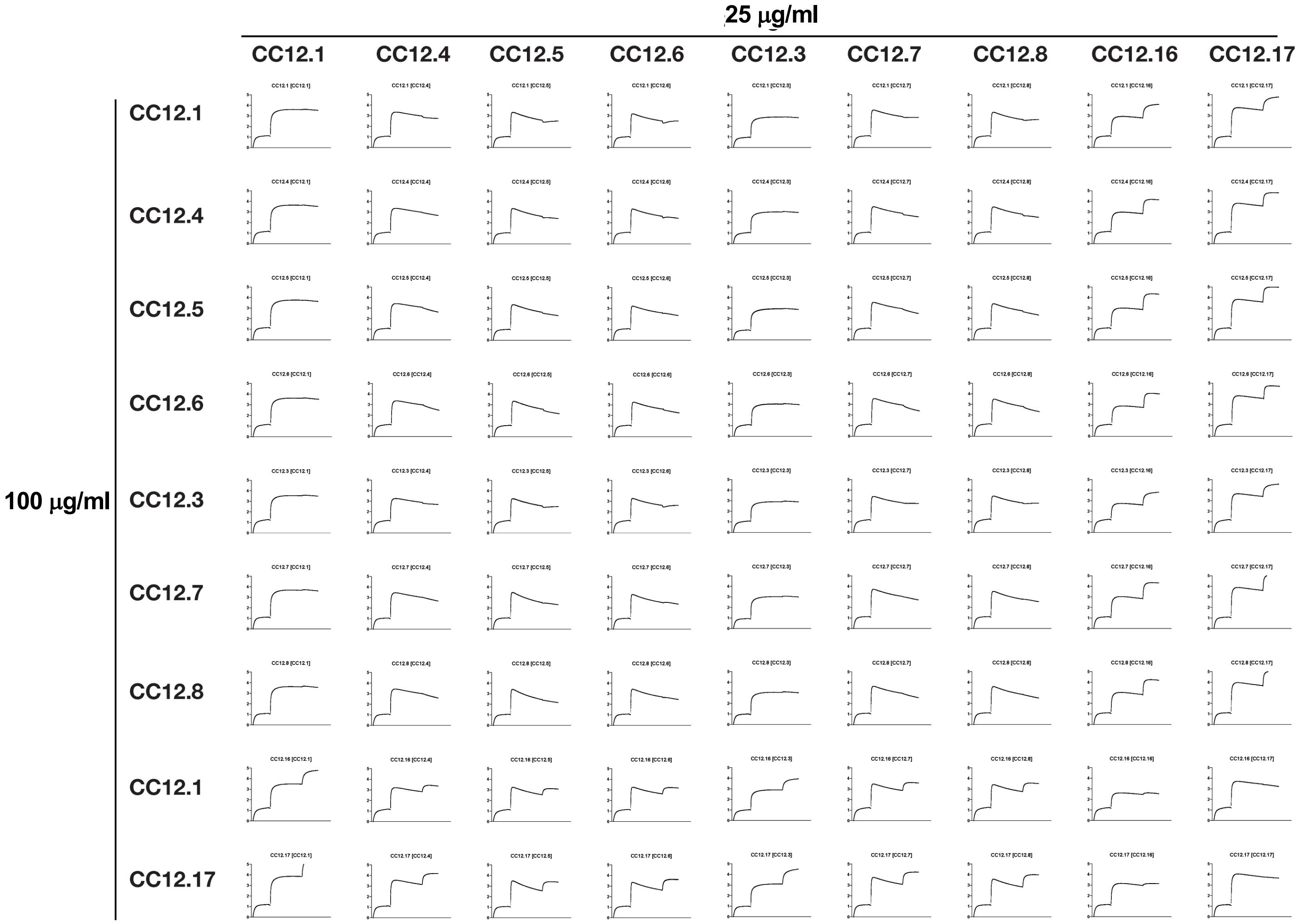

**Fig. S7A: Monoclonal antibodies were evaluated for epitope competition using an Octet RED384 platform**. His-tagged RBD was captured using an anti-Penta-HIS biosensor and indicated mAbs at a concentration of 100 µg/ml are first incubated for 10 min followed by incubation with 25 µg/ml of competing antibodies for 5 min.

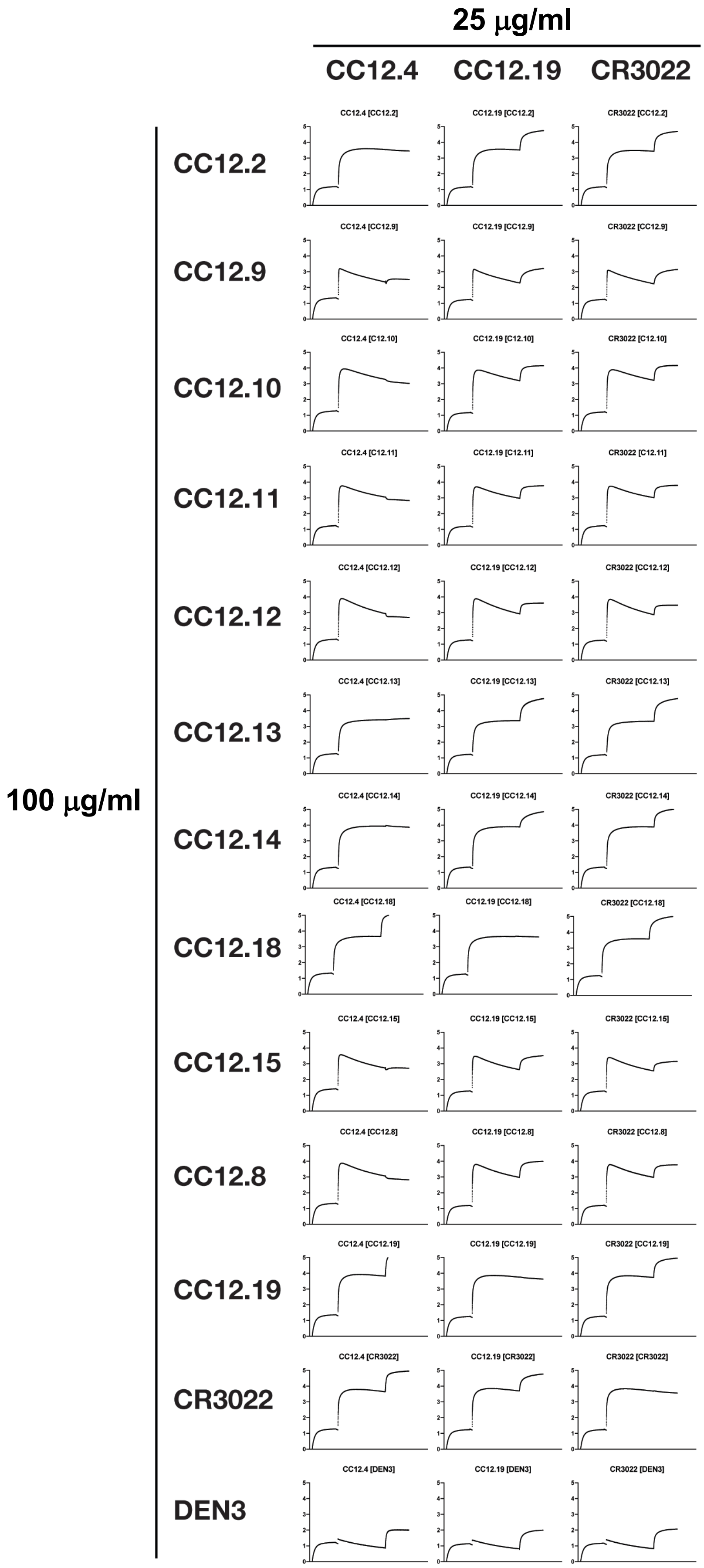

**Fig. S7B: Monoclonal antibodies were evaluated for epitope competition using an Octet RED384 platform**. His-tagged RBD was captured using an anti-Penta-HIS biosensor and indicated mAbs at a concentration of 100 μg/ml are first incubated for 10 min followed by incubation with 25 μg/ml of competing antibodies for 5 min.

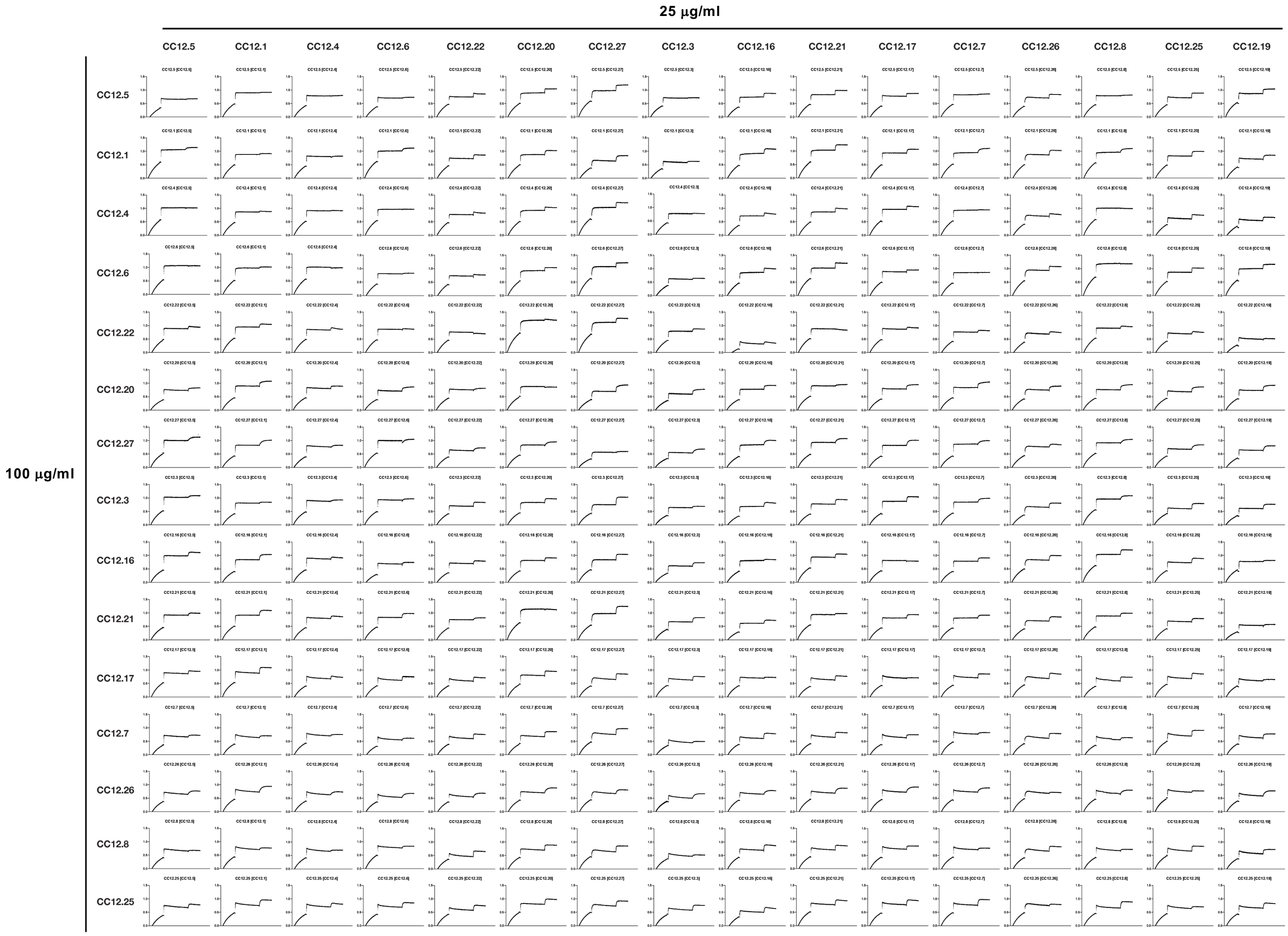

**Fig. S7C: Monoclonal antibodies were evaluated for epitope competition using an Octet RED384 platform**. His-tagged RBD was captured using an anti-Penta-HIS biosensor and indicated mAbs at a concentration of 100 μg/ml are first incubated for 10 min followed by incubation with 25 μg/ml of competing antibodies for 5 min.

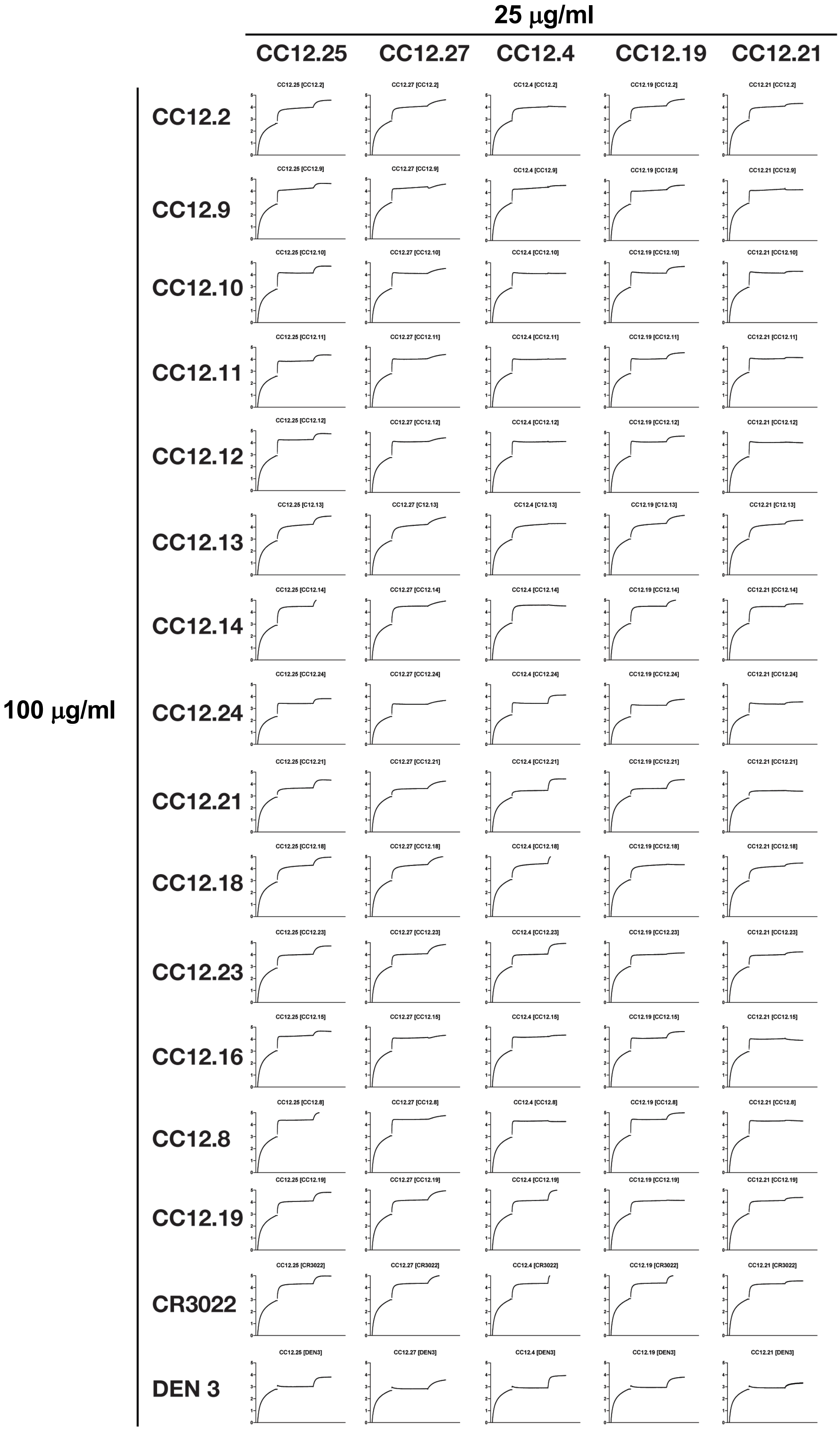

**Fig. S7D: Monoclonal antibodies were evaluated for epitope competition using an Octet RED384 platform**. His-tagged RBD was captured using an anti-Penta-HIS biosensor and indicated mAbs at a concentration of 100 μg/ml are first incubated for 10 min followed by incubation with 25 μg/ml of competing antibodies for 5 min.

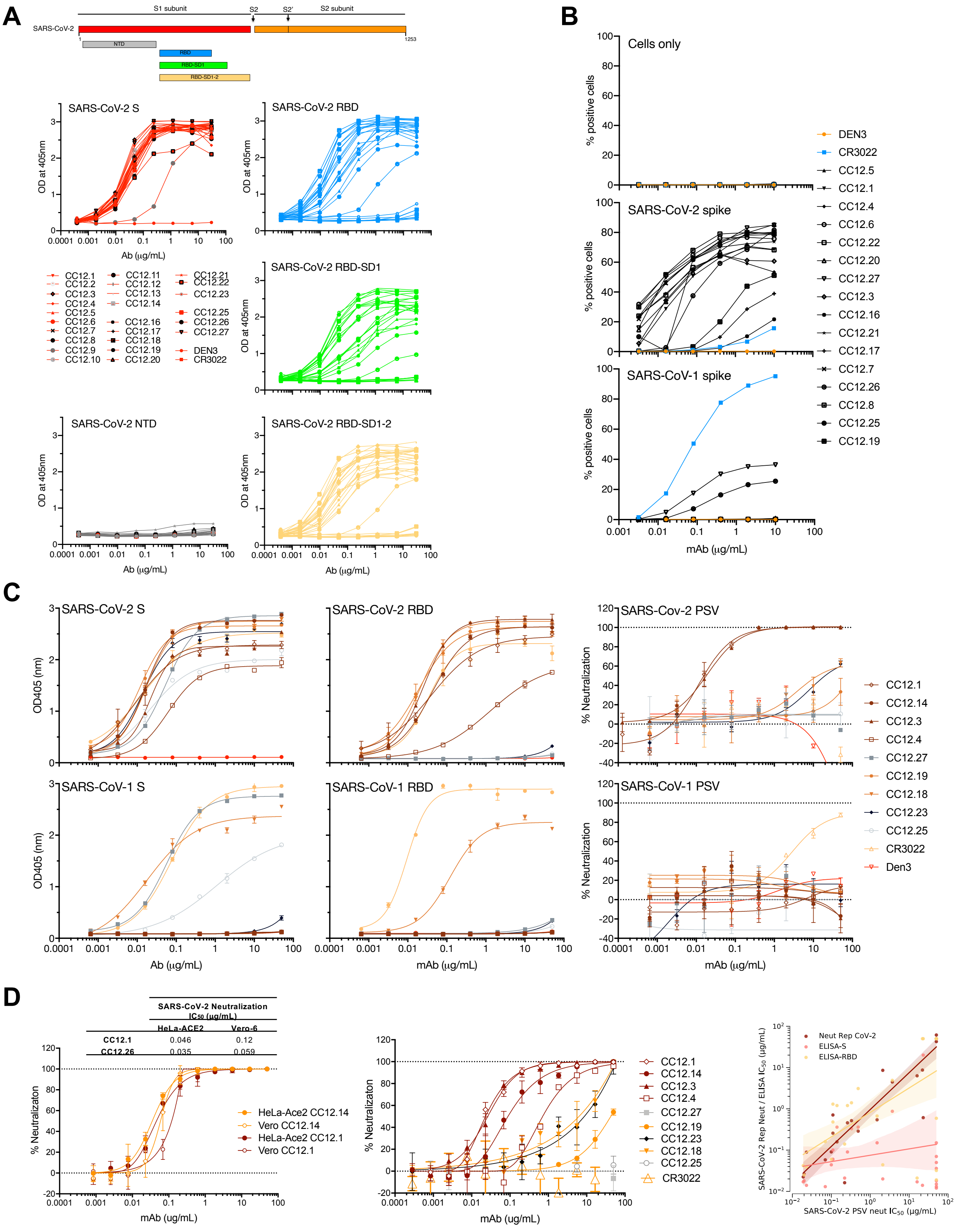

**Fig S8. Functional characterization of SARS-CoV-2 specific mAbs selected from the HTP screening.** Antibodies were tested for (**A**) ELISA for binding to several truncated versions of the recombinant SARS-CoV-1 and SARS-CoV-2 S proteins, (**B**) cell surface binding to SARS-CoV-1 and CoV-2 Spike, and for neutralization of (**C**) pseudotyped and (**D**) replicating SARS-CoV-2 on HeLa-ACE2 or Vero (middle plot only). **(E)** ACE2 binding inhibition was performed by flow cytometry. Purified mAbs were mixed with biotinylated SARS-CoV-2 S or RBD proteins at a molar ratio of 4:1 before adding to HeLa-ACE2 cells. Streptavidin-AF647 was used to detect the relative intensity of cell surface ACE2 binding biotinylated SARS-CoV-2 S or RBD proteins. SARS-CoV-2 S or RBD-AF647 mean fluorescence intensity (MFI) was determined from the gate of singlet and PI negative cells. CR3022 and Den3 were used as negative controls. HeLa and HeLa-ACE2 cells stained with SARS-CoV-2 S or RBD alone were used as background and positive control separately.

**E**

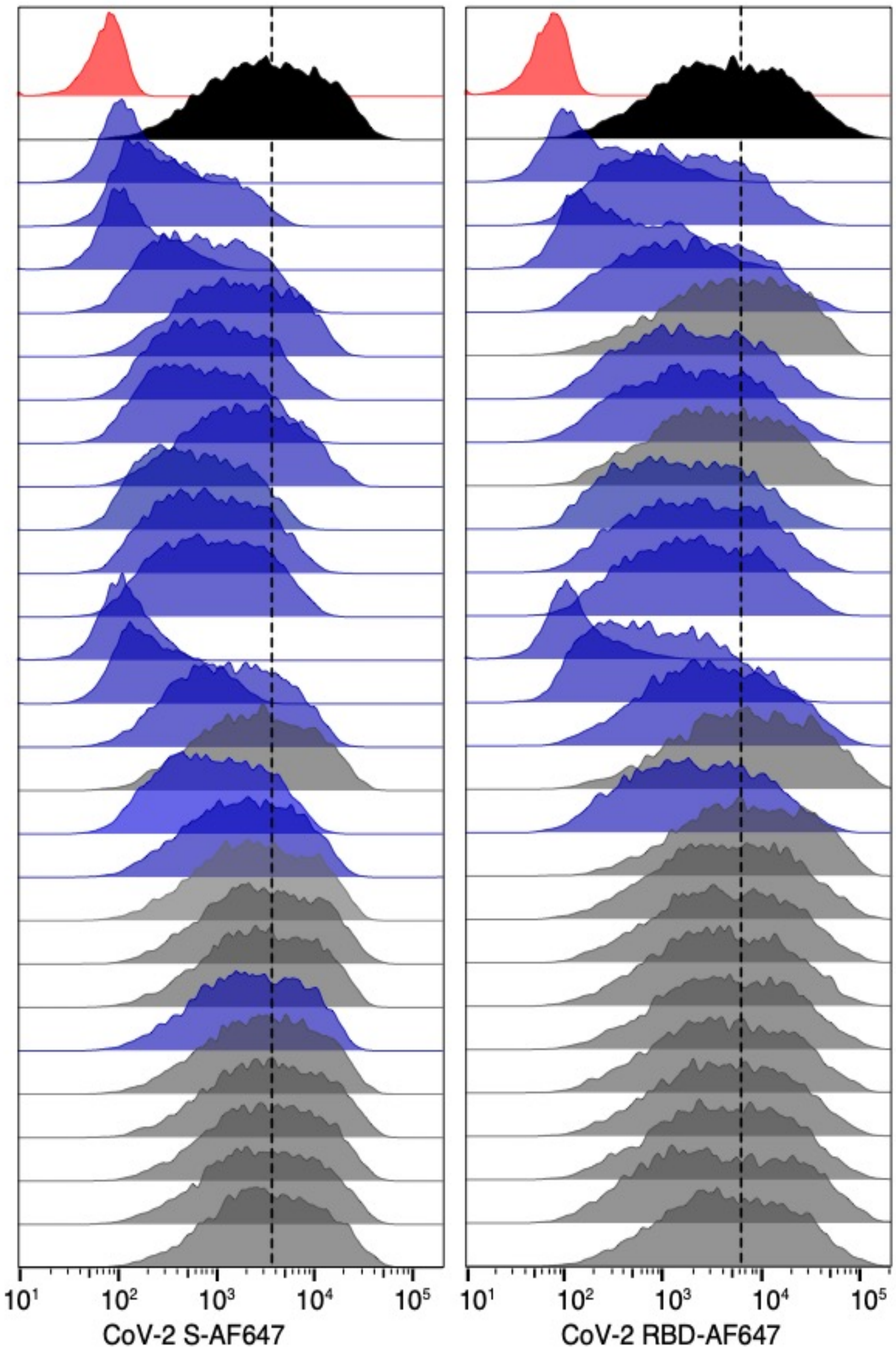

| mAbs | MFI of CoV-2 S | MFI of CoV-2 RBD |
|---|---|---|
| Background | 80.0 | 74.5 |
| Probe alone | 6502 | 10817 |
| CC12.1 | 165 | 528 |
| CC12.2 | 731 | 3682 |
| CC12.3 | 214 | 982 |
| CC12.4 | 1430 | 5194 |
| CC12.5 | 3155 | 11672 |
| CC12.6 | 1756 | 5167 |
| CC12.8 | 1369 | 5105 |
| CC12.9 | 3765 | 9714 |
| CC12.10 | 1211 | 4036 |
| CC12.11 | 1681 | 5614 |
| CC12.12 | 1787 | 5738 |
| CC12.13 | 145 | 178 |
| CC12.14 | 463 | 2228 |
| CC12.15 | 2736 | 8718 |
| CC12.16 | 4945 | 18054 |
| CC12.18 | 1792 | 5711 |
| CC12.19 | 3726 | 16358 |
| CC12.20 | 4959 | 11677 |
| CC12.21 | 5869 | 11140 |
| CC12.22 | 5504 | 10020 |
| CC12.23 | 4189 | 11601 |
| CC12.25 | 5834 | 11207 |
| CC12.26 | 6660 | 11004 |
| CC12.27 | 6219 | 10918 |
| CR3022 | 6077 | 12763 |
| Den3 | 6567 | 11681 |

**Fig S8. Functional characterization of SARS-CoV-2 specific mAbs selected from the HTP screening.** Antibodies were tested for (**A**) ELISA for binding to several truncated versions of the recombinant SARS-CoV-1 and SARS-CoV-2 S protein, (**B**) cell surface binding to SARS-CoV-1 and CoV-2 Spike, and for neutralization of (**C**) pseudotyped and (**D**) replicating SARS-CoV-2 on HeLa-ACE2 or Vero (middle plot only). **(E)** ACE2 binding inhibition was performed by flow cytometry. Purified mAbs were mixed with biotinylated SARS-CoV-2 S or RBD proteins at a molar ratio of 4:1 before adding to HeLa-ACE2 cells. Streptavidin-AF647 was used to detect the relative intensity of cell surface ACE2 binding biotinylated SARS-CoV-2 S or RBD proteins. SARS-CoV-2 S or RBD-AF647 mean fluorescence intensity (MFI) was determined from the gate of singlet and PI negative cells. CR3022 and Den3 were used as negative control. HeLa and HeLa-ACE2 cells stained with SARS-CoV-2 S or RBD alone were used as background and positive control separately.

**A**

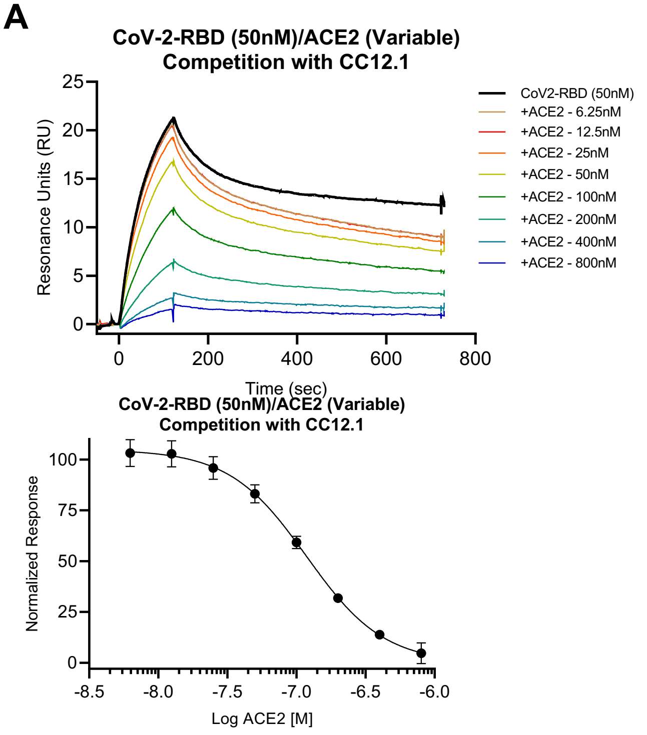

**B**

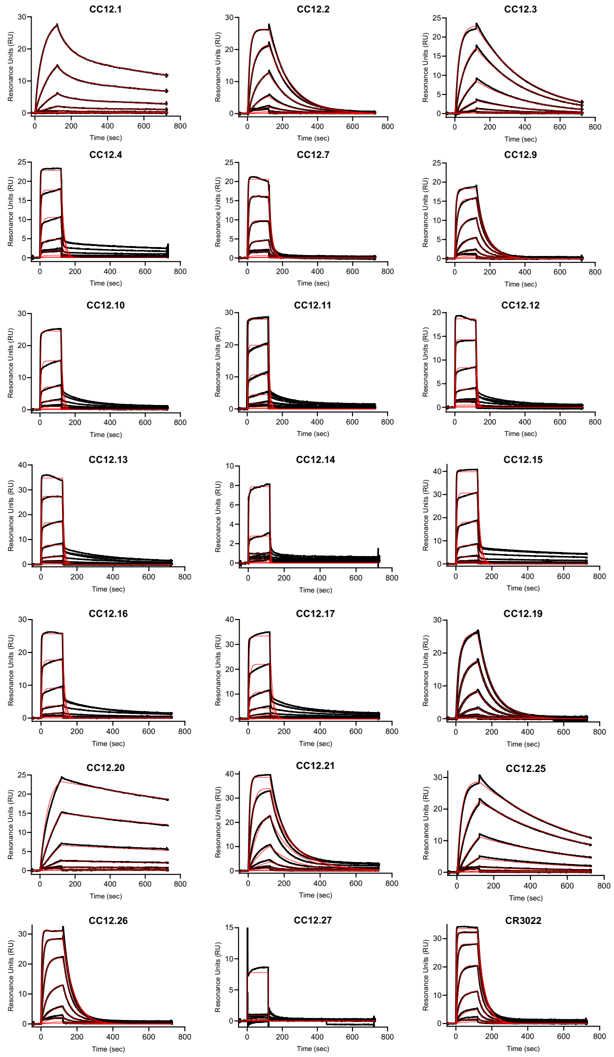

**C**

| mAb ID | Method | Analyte | Fit Model | ka [1/Ms] | kd [1/s] | KD [M] |
|---|---|---|---|---|---|---|
| CC12.1 | Fc-Capture | CoV-2-RBD | 1:1 Binding | 6.95E+05 | 4.12E-03 | 5.92E-09 |
| CC12.2 | Fc-Capture | CoV-2-RBD | 1:1 Binding | 1.48E+05 | 8.03E-03 | 5.43E-08 |
| CC12.3 | Fc-Capture | CoV-2-RBD | 1:1 Binding | 3.99E+05 | 3.42E-03 | 8.59E-09 |
| CC12.4 | Fc-Capture | CoV-2-RBD | 1:1 Binding | 3.88E+05 | 6.60E-02 | 1.70E-07 |
| CC12.5 | Fc-Capture | CoV-2-RBD | 1:1 Binding | - | - | ND |
| CC12.6 | Fc-Capture | CoV-2-RBD | 1:1 Binding | - | - | ND |
| CC12.7 | Fc-Capture | CoV-2-RBD | 1:1 Binding | 3.80E+05 | 6.03E-02 | 1.59E-07 |
| CC12.8 | Fc-Capture | CoV-2-RBD | 1:1 Binding | - | - | ND |
| CC12.9 | Fc-Capture | CoV-2-RBD | 1:1 Binding | 3.64E+05 | 1.70E-02 | 4.66E-08 |
| CC12.10 | Fc-Capture | CoV-2-RBD | 1:1 Binding | 4.36E+05 | 9.61E-02 | 2.20E-07 |
| CC12.11 | Fc-Capture | CoV-2-RBD | 1:1 Binding | 4.93E+05 | 1.26E-01 | 2.55E-07 |
| CC12.12 | Fc-Capture | CoV-2-RBD | 1:1 Binding | 7.82E+05 | 1.43E-01 | 1.83E-07 |
| CC12.13 | Fc-Capture | CoV-2-RBD | 1:1 Binding | 7.73E+05 | 1.20E-01 | 1.55E-07 |
| CC12.14 | Fc-Capture | CoV-2-RBD | 1:1 Binding | 5.94E+04 | 3.12E-01 | >5.00E-07 |
| CC12.15 | Fc-Capture | CoV-2-RBD | 1:1 Binding | 7.86E+05 | 6.91E-02 | 8.79E-08 |
| CC12.16 | Fc-Capture | CoV-2-RBD | 1:1 Binding | 5.75E+05 | 8.43E-02 | 1.47E-07 |
| CC12.17 | Fc-Capture | CoV-2-RBD | 1:1 Binding | 4.04E+05 | 7.07E-02 | 1.75E-07 |
| CC12.18 | Fc-Capture | CoV-2-RBD | 1:1 Binding | - | - | ND |
| CC12.19 | Fc-Capture | CoV-2-RBD | 1:1 Binding | 4.43E+05 | 1.15E-02 | 2.60E-08 |
| CC12.20 | Fc-Capture | CoV-2-RBD | 1:1 Binding | 1.40E+05 | 3.64E-04 | 2.60E-09 |
| CC12.21 | Fc-Capture | CoV-2-RBD | 1:1 Binding | 4.35E+05 | 1.57E-02 | 3.62E-08 |
| CC12.22 | Fc-Capture | CoV-2-RBD | 1:1 Binding | - | - | ND |
| CC12.23 | Fc-Capture | CoV-2-RBD | 1:1 Binding | - | - | ND |
| CC12.24 | Fc-Capture | CoV-2-RBD | 1:1 Binding | - | - | ND |
| CC12.25 | Fc-Capture | CoV-2-RBD | 1:1 Binding | 4.28E+05 | 1.66E-03 | 3.88E-09 |
| CC12.26 | Fc-Capture | CoV-2-RBD | 1:1 Binding | 6.36E+05 | 1.70E-02 | 2.68E-08 |
| CC12.27 | Fc-Capture | CoV-2-RBD | 1:1 Binding | 4.45E+03 | 1.64E+00 | >1.00E-6 |
| CR3022 | Fc-Capture | CoV-2-RBD | 1:1 Binding | 1.03E+06 | 2.54E-02 | 2.47E-08 |

**Fig. S9: SARS-CoV-2 nAb affinities.** (**A**) CC12.1 binding inhibits the interaction of SARS-CoV-2-RBD with ACE2. (Top) Representative data from a SPR competition experiment in which the effect of varying concentrations of ACE2 on the interaction of 50 nM SARS-CoV-2-RBD with CC12.1 was examined. A legend showing the identity of each sensorgram is *inset*, in which the ACE2 injection alone was subtracted from the SARS-CoV-2-RBD + ACE2 injection series. The residual SARS-CoV-2-RBD binding in the presence of multiple ACE2 concentrations is shown in varying colors, while the sensorgram for the same concentration of SARS-CoV-2-RBD in the absence of ACE2 is shown as the darkest black line. (Bottom) Residual SARS-CoV-2-RBD binding in the presence of various concentrations of ACE2, normalized to 50 nM CoV-2-RBD (100%) and buffer (0%), and fit to a dose-response curve ($IC_{50}$ = 120 nM). (**B**) SPR sensorgrams for SARS-CoV-2-RBD binding. Recombinant antibodies were captured via Fc-capture to an anti-human IgG (Fc) antibody and varying concentrations of SARS-CoV-2-RBD were injected using a multi-cycle method. Representative sensorgrams in resonance units (RUs) plotted against time of injection are shown. Black lines are the experimental trace obtained from the SPR experiments and red are the best global fits (1:1 Langmuir binding model) to the data used to calculate the association (ka) and dissociation (kd) rate constants. (**C**) Summarized results of SARS-CoV-2-RBD binding to antibodies via a Fc-capture, multi-cycle method. Association and dissociation rate constants calculated through a 1:1 Langmuir binding model using the BIAevaluation software. ND = not determined.

| mAb name | Predicted Unique Epitope | ELISA $IC_{50}$ (µg/mL) | | | | Poly-reactivity | PSV Neutralization $IC_{50}$ (µg/mL) | | | Live virus SARS-CoV-2 |
|---|---|---|---|---|---|---|---|---|---|---|
| | | SARS-CoV-2 S | SARS-CoV-2 RBD | SARS-CoV-1 S | SARS-CoV-1 RBD | | SARS-CoV-1 | SARS-CoV-2 | MPN | |
| **CC12.1** | RBD-A | 0.017 | 0.042 | >50 | >50 | None | >50 | 0.019 | 100% | 0.022 |
| **CC12.2** | RBD-A | 0.021 | 0.14 | >50 | >50 | None | >50 | 0.22 | 100% | 0.016 |
| **CC12.3** | RBD-A | 0.018 | 0.025 | >50 | >50 | None | >50 | 0.018 | 100% | 0.026 |
| **CC12.4** | RBD-A | 0.062 | 1.3 | >50 | >50 | None | >50 | 0.11 | 100% | 0.71 |
| **CC12.5** | RBD-A | 0.33 | 1.4 | >50 | >50 | None | >50 | 0.33 | 100% | 0.52 |
| **CC12.6** | RBD-A | 0.36 | 3.1 | >50 | >50 | None | >50 | 0.49 | 100% | 0.29 |
| **CC12.7** | RBD-A | 0.091 | 1.2 | >50 | >50 | None | >50 | 0.26 | 89% | 0.46 |
| **CC12.8** | RBD-A | 0.10 | 0.48 | >50 | >50 | None | >50 | 0.11 | 100% | 0.26 |
| **CC12.9** | RBD-A | 20 | >50 | >50 | >50 | None | >50 | 23 | 60% | 0.61 |
| **CC12.10** | RBD-A | 0.013 | 0.16 | >50 | >50 | None | >50 | 0.07 | 95% | 0.058 |
| **CC12.11** | RBD-A | 0.36 | 1.2 | >50 | >50 | None | >50 | 0.14 | 85% | 0.16 |
| **CC12.12** | RBD-A | 0.15 | 0.67 | >50 | >50 | ND | >50 | 1.5 | 99% | 0.54 |
| **CC12.13** | RBD-A | 0.044 | 0.058 | >50 | >50 | None | >50 | 0.10 | 100% | 0.090 |
| **CC12.14** | RBD-A | 0.014 | 0.039 | >50 | >50 | ND | >50 | 0.023 | 100% | 0.089 |
| **CC12.15** | RBD-A | 0.87 | 4.9 | >50 | >50 | None | >50 | 3.7 | 65% | 4.5 |
| **CC12.16** | RBD-B | 0.073 | 0.59 | >50 | >50 | None | >50 | >50 | 21% | 61 |
| **CC12.17** | RBD-B | 0.020 | 0.044 | >50 | >50 | None | >50 | 2.1 | 91% | 8.6 |
| **CC12.18** | RBD-B | 0.017 | 0.021 | 0.018 | 2.1 | None | >50 | 16 | 66% | 6.3 |
| **CC12.19** | RBD-B | 0.012 | 0.019 | >50 | >50 | None | >50 | >50 | 50% | 43 |
| **CC12.20** | SPIKE-A | 0.036 | >50 | >50 | >50 | None | >50 | >50 | | >50 |
| **CC12.21** | SPIKE-A | 0.0050 | >50 | >50 | >50 | None | >50 | >50 | | >50 |
| **CC12.22** | SPIKE-A | >50 | >50 | >50 | >50 | None | >50 | >50 | 41% | >50 |
| **CC12.23** | SPIKE-A | 0.014 | >50 | >50 | >50 | None | >50 | 22 | 64% | 8.7 |
| **CC12.24** | ND | 0.013 | >50 | >50 | >50 | None | >50 | >50 | | ND |
| **CC12.25** | SPIKE-B | 0.018 | >50 | 1.1 | >50 | None | >50 | >50 | | >50 |
| **CC12.26** | SPIKE-B | 0.15 | >50 | >50 | >50 | None | >50 | >50 | | >50 |
| **CC12.27** | SPIKE-C | 0.053 | >50 | 0.057 | >50 | None | >50 | >50 | | >50 |
| **CR3022** | RBD-C | 0.015 | 0.032 | 2.9 | 2.7 | None | 3.0 | >50 | | |
| **Den3** | | >50 | >50 | >50 | >50 | None | >50 | >50 | | |

**Fig. S10: SARS-CoV-2 nAb functional summary.** SARS-Cov-1 and SARS-CoV-2 binding affinities and neutralization potency of the indicated SARS-CoV-2 specific mAbs isolated from CC12 are tabulated. Neutralization was tested against pseudotyped (PSV) and live replicating SARS-CoV-1 and SARS-CoV-2 viruses. MPN: Maximum Neutralization Plateau.

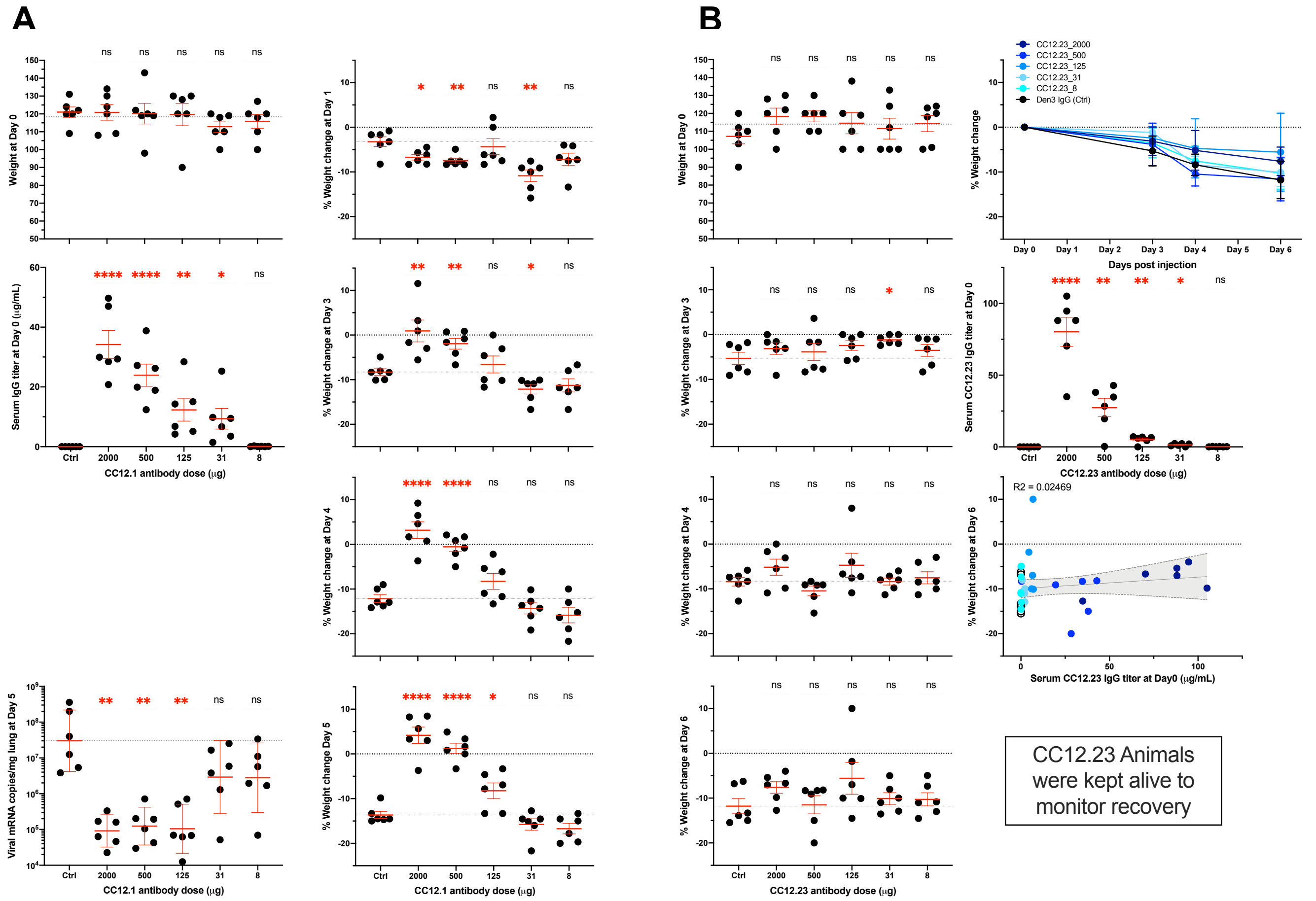

**Fig.S11: Animal protection studies.** Syrian hamsters received various doses of SARS-CoV-2-specific human mAbs CC12.1 or CC12.23 or 2 mg dengue-specific human mAb Den3 i.p 12h before i.n. challenge with SARS-CoV-2. (**A**) CC12.1 protection experiment. From top to bottom, left to right: Weights of animals at time of challenge (Day 0); CC12.1 serum concentration in each animal as measured by ELISA at time of challenge (12h post administration, Day 0); Viral load in lung tissue for each animal at day5-post challenge; linear correlation between serum human IgG concentration at time of termination (Day 5) and % weight loss at day 5 (95% confidence intervals indicated in grey shade, R-square value is also indicated); Weight change (%) in each animal at Day 1,3,4 and 5 post challenge. (**B**) CC12.23 protection experiment. From top to bottom, left to right: Weights of animals at time of challenge (Day 0); Weight change (%) in each animal at Day 3, 4 and 5 post challenge (Day 0); average weight loss in each group over time.; linear correlation between serum human IgG concentration at time of challenge (Day 0) and % weight loss at Day 5 (95% confidence intervals indicated in grey shade, R-square value is also indicated). Significance of the difference between the groups was evaluated with Mann-Whitney U-tests using a 95% confidence interval. P-values are indicated (ns: non-significant; (*) <0.0332; (**) <0.0021; (***) <0.0002; (****) <0.00001).

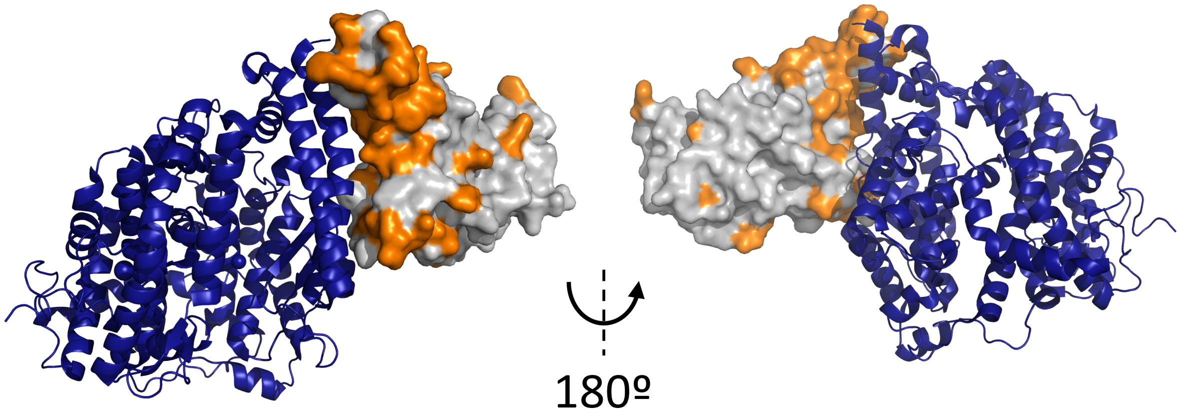

**Fig.S12: Crystal structure of SARS-CoV-2 RBD bound to human ACE2 (adapted from PDBID 6M0J)**. ACE2 is represented as blue ribbons and RBD is surface rendered, with mutations from SARS-CoV-1 highlighted in orange.
